# NK-like and networked CD8^+^ T cell immunity mediates exceptional HIV control

**DOI:** 10.64898/2026.07.02.735504

**Authors:** Aljawharah Alrubayyi, Anna K. Traunbauer, Charles R. Crain, Shaown Bhattacharyya, David R. Collins, Hsinyen Huang, Clarety Kaseke, Umar Arshad, Xiaodong Lian, Aarthi Vijayakumar, Matthew A. Getz, Mathias Lichterfeld, Xu G. Yu, Bruce D. Walker, Georg M. Lauer, Gaurav D. Gaiha

## Abstract

Durable treatment-free remission remains a defining goal for people living with HIV (PLWH). Studies of spontaneous elite controllers have revealed that functional CD8⁺ T cells targeting structurally networked viral epitopes can mediate durable viral suppression^1,2^. However, rare reservoir-defined exceptional controllers within the spectrum of elite control^3–5^, characterized by the absence of intact provirus or proviruses confined to transcriptionally repressed genomic regions^6^, provide a unique opportunity to define mechanisms of cure-like immunity. Here, we integrate functional epitope mapping, single-cell transcriptomics, and infected cell elimination assays to identify networked HIV epitope targeting and a natural killer (NK)-like killer-cell immunoglobulin-like receptor (KIR)⁺ CD8⁺ T cell subset as key features of exceptional control. This NK-like subset was selectively enriched within HIV-specific, but not CMV-specific, CD8⁺ T cells from controllers, and was transcriptionally similar to highly cytotoxic subsets within the broader KIR^+^ CD8^+^ T cell compartment. Flow cytometry revealed increased frequencies of KIR⁺ CD8⁺ T cells in exceptional controllers relative to antiretroviral therapy (ART)-suppressed individuals, and unexpectedly, enrichment of dual KIR^+^ NKG2A^+^ CD8⁺ T cells. Functional depletion of KIR⁺ CD8⁺ T cells significantly impaired the elimination of autologous HIV-infected CD4⁺ T cells, despite preserved recognition by proliferative networked HIV-specific CD8⁺ T cells. These findings thereby identify an NK-like KIR⁺ CD8⁺ T cell state as a previously unrecognized component of exceptional HIV immunity that complements networked epitope targeting, providing a novel framework for immunotherapeutic HIV cure strategies.

## Main Text

Spontaneous elite controllers (ECs), who comprise less than 1% of PLWH, demonstrate that durable viral suppression can occur without antiretroviral therapy (ART) and have provided a critical model for defining immune mechanisms of natural control^7^. Prior studies have linked this phenotype to specific HLA class I gene polymorphisms^8–10^, proliferative HIV-specific CD8⁺ T cells^11–13^, lytic granule loading^14^, and preferential targeting of mutationally constrained, structurally networked viral epitopes that limit immune escape^1,2,15^. These observations support a model in which effective HIV control depends on the coupling of CD8⁺ T cell functional capacity with recognition of epitopes derived from viral sites that are difficult for HIV to mutate without compromising fitness.

Within this spectrum of spontaneous elite control, rarer exceptional elite controller (EEC) individuals exhibit a cure-like reservoir phenotype^3–5^, in which intact proviruses are undetectable (type I cure; EEC-I) or confined to deeply transcriptionally repressed genomic regions (type II cure; EEC-II), such as centromeric satellite DNA and KRAB-zinc finger loci^6^. These EEC-I and EEC-II individuals represent the extreme end of spontaneous elite control and provide a unique opportunity to delineate immune features associated with an HIV cure^16^. Notably, one EEC-I, the late San Francisco patient^17,18^, was shown to lack detectable intact provirus across extensive sampling of blood and tissue^3^, with intact reservoir measurements lower than those observed in the cured Berlin patient^6^, who underwent stem cell transplantation with homozygous CCR5Δ32 donor cells^19^. Although such observations do not prove sterilizing cure, they indicate that near-complete immune-mediated clearance or durable silencing of intact provirus may be achievable. Defining the immune features associated with this state could therefore reveal cellular programs that therapeutic vaccination or immunotherapy may need to elicit to achieve sustained HIV remission.

Here, we evaluate CD8⁺ T cell responses in reservoir-defined EECs and ART-suppressed PLWH using functional epitope mapping, autologous viral sequence analysis, single-cell transcriptomics, surface marker assessments, and functional elimination assays of autologous HIV-infected CD4⁺ T cells. This approach identifies networked epitope targeting by proliferative CD8^+^ T cell responses as a feature of exceptional control. However, unexpectedly, our studies revealed the enrichment of a highly cytotoxic NK-like subset within EEC HIV-specific CD8⁺ T cells. This subset contributes non-redundantly to the elimination of autologous HIV-infected CD4^+^ T cells and shares striking transcriptional similarity with virus-specific CD8^+^ T cells observed in functional HBV cure (Lattouf et al.; co-submitted manuscript). These findings thereby demonstrate that an NK-like CD8⁺ T cell program complements networked epitope targeting in exceptional HIV control.

### Exceptional controllers preferentially target networked HIV epitopes with proliferative CD8^+^ T cell responses

To define the specificity and functional capacity of HIV-specific CD8⁺ T cell responses in exceptional control, we performed IFN-γ ELISpot and CFSE-based proliferation assays using HLA-matched optimal HIV clade B epitopes in EECs (n = 10) and ART-suppressed individuals (n = 22) (**Fig. 1a; Table 1; Extended Data Fig. 1**). The two groups were comparable in age, sex, race, and viral suppression, but as expected, EECs were enriched for protective HLA class I alleles^8^ and the compound KIR3DL1^+^/HLA-Bw4^+^ genotype that is associated with improved outcomes^20^ (**Table 1, Extended Table 1**). Notably, individual inhibitory KIR genes (*KIR2DL1, KIR2DL3, KIR3DL1*) did not distinguish the two groups. We first evaluated T cell responses to all HLA-matched epitopes for each individual and found that the summed magnitude and breadth of *ex vivo* IFN-γ responses did not differ significantly between EECs and ART-suppressed individuals (**Fig. 1b, c**). However, when the analysis was restricted to mutationally constrained networked epitopes, which were restricted by both protective (*B*57, B*27*) and non-protective (*A*11, B*07*) HLA class I alleles, EECs exhibited greater IFN-γ response magnitude and breadth than ART-suppressed individuals (**Fig. 1d, e**). Moreover, immunodominant responses in exceptional controllers (defined for each participant as the response with the highest *ex vivo* IFN-γ ELISpot magnitude) had significantly higher epitope network scores (**Fig. 1f**), indicating that exceptional control is associated with preferential targeting of structurally networked viral regions rather than a generalized increase in HIV-specific T cell reactivity.

**Table 1.** Clinical and immunogenetic characteristics of study participants. Statistical comparisons: continuous variables, two-sided Mann-Whitney U test; categorical variables, two-sided Fisher’s exact test. Significant P values (<0.05) shown in bold. EEC, exceptional elite controller; ART, antiretroviral therapy-suppressed. *a* Protective HLA allele defined as expression of ≥1 of HLA-B*57, B*52, B*14:02, or B*27:05, or HLA-A*25, as associated with HIV control in prior immunogenetic studies8. Individual allele assignments are provided in the Extended Table. *b* Compound KIR3DL1/HLA-Bw4 genotype, the protective inhibitory receptor-ligand pairing associated with HIV control^2^0.

| Characteristic | EEC (n = 10) | ART-suppressed (n = 22) | P-value |
| --- | --- | --- | --- |
| Demographic and clinical |  |  |  |
| Age, years - median (range) | 68 (58-80) | 66 (43-82) | 0.611 |
| Female sex - n (%) | 2 (20) | 4 (18) | 1.000 |
| White - n (%) | 8 (80) | 17 (77) | 1.000 |
| Black or African American - n (%) | 2 (20) | 5 (23) | 1.000 |
| CD4<sup>+</sup> T cells, cells/μl - median (range) | 808 (450-2168) | 642 (305-1456) | **0.033** |
| Plasma HIV-1 RNA, copies/ml | <20 | <20 | 1.000 |
| HLA class I genotype |  |  |  |
| Protective HLA allele - n (%)<sup>a</sup> | 9 (90) | 6 (27) | **0.002** |
| HLA-Bw4 motif - n (%) | 9 (90) | 12 (55) | 0.106 |
| KIR3DL1<sup>+</sup>/HLA-Bw4<sup>+</sup> - n (%)<sup>b</sup> | 9 (90) | 10 (45) | **0.024** |
| KIR genotype |  |  |  |
| KIR2DL1 - n (%) | 10 (100) | 21 (100) | 1.000 |
| KIR2DL3 - n (%) | 10 (100) | 20 (95) | 1.000 |
| KIR3DL1 - n (%) | 10 (100) | 20 (95) | 1.000 |

**Figure 1.**
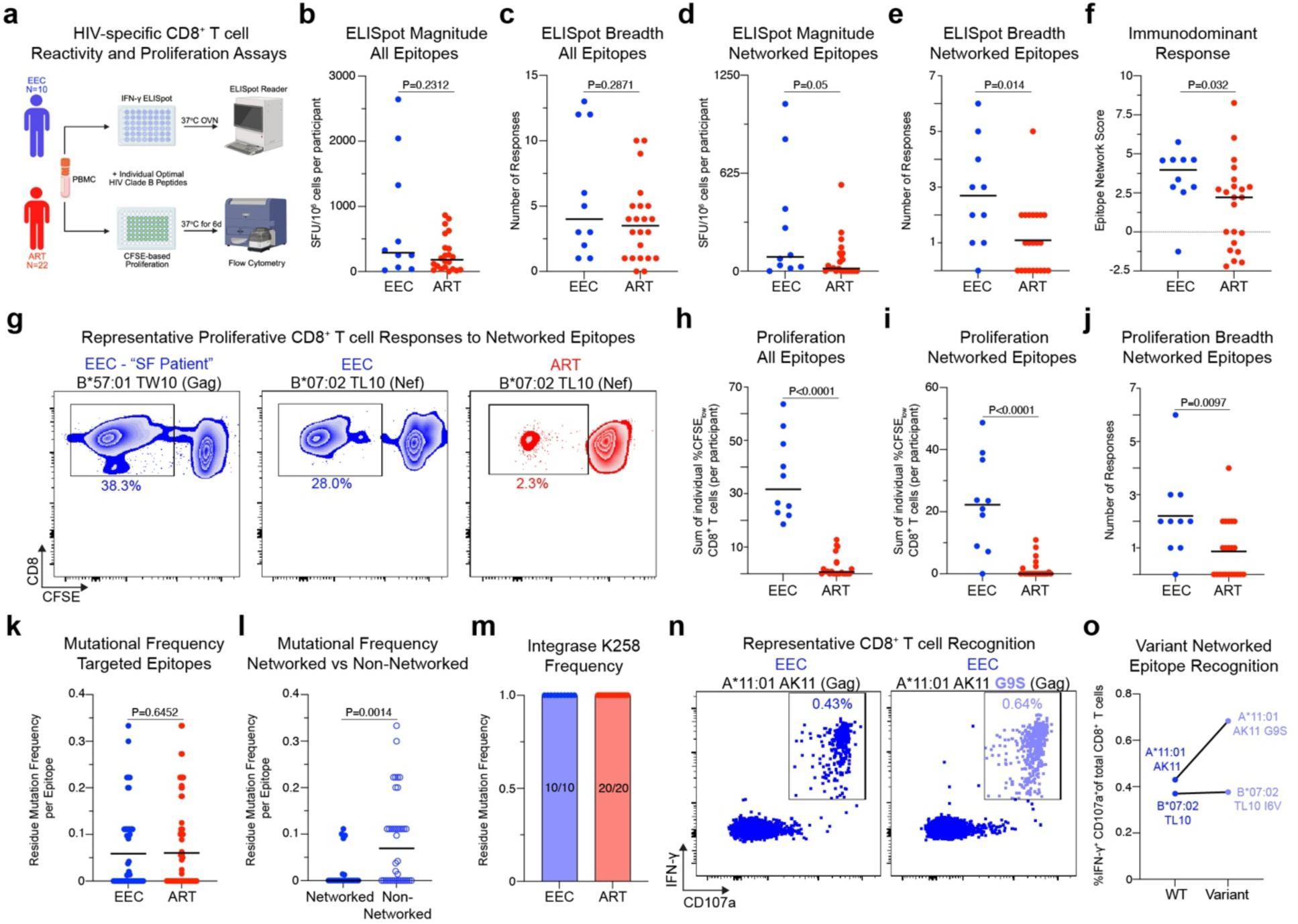
Targeting of structurally networked epitopes with proliferative CD8^+^ T cell responses distinguishes exceptional controllers. **a**, Schematic overview of the HIV-specific CD8⁺ T cell reactivity and proliferation assay workflow performed in exceptional elite controllers (EECs; blue, n = 10) and ART-suppressed individuals (ART; red, n = 22). PBMCs were stimulated with HLA-matched optimal HIV clade B epitopes and assessed by IFN-γ ELISpot and CFSE-based proliferation assays. **b, c**, Total magnitude (b) and breadth (c) of IFN-γ ELISpot responses to all HLA-matched optimal epitopes. Magnitude is shown as spot-forming units (SFU) per 10⁶ PBMCs; breadth is the number of positive responses (>10 SFU per 10⁶ PBMCs). **d, e**, Magnitude (d) and breadth (e) of IFN-γ ELISpot responses restricted to networked epitopes. **f**, Epitope network scores of immunodominant CD8⁺ T cell responses in EECs and ART individuals. Each point represents the immunodominant response per participant; dashed line indicates a network score of zero. **g**, Representative CFSE-based proliferative CD8⁺ T cell responses to networked epitopes in one EEC-I, one EEC-II, and one ART-treated individual. **h**, Summed magnitude of proliferative CD8⁺ T cell responses (%CFSE_low_ CD8⁺ T cells) to all HLA-matched optimal epitopes per participant. **i, j**, Magnitude (i) and breadth (j) of proliferative CD8⁺ T cell responses to networked epitopes. Breadth is the number of individual networked epitopes with positive proliferative responses. **k**, Residue mutation frequency within targeted optimal HIV epitopes in EECs and ART individuals, calculated as the percentage of proviral sequence clones containing at least one amino acid difference from the HIV clade B consensus sequence. **l**, Residue mutation frequency in networked versus non-networked epitopes within EECs. **m**, Frequency of the HIV integrase K258 residue in EECs and ART individuals. Numbers indicate the number of participants with the K258 residue out of the total number analyzed. **n**, Representative flow cytometry plots showing CD8⁺ T cell recognition of wild-type and variant (G9S) A11:01-restricted AK11 Gag epitopes, assessed by intracellular IFN-γ and CD107a expression. **o**, Percentage of IFN-γ⁺CD107a⁺ CD8⁺ T cell responses to wild-type and variant networked epitopes across participants. Each dot represents one epitope-specific response in one participant. Statistical comparisons in **b-f, h-l** were performed using non-parametric Mann-Whitney *U*-tests.

We next assessed the proliferative capacity of HIV-specific CD8^+^ T cell responses, given the established link to spontaneous elite HIV control^11–13^. In contrast to the modest differences observed by ELISpot, analysis of all HLA-matched epitopes revealed markedly higher proliferative responses (%CD8^+^ CFSE_low_) in exceptional controllers relative to ART-suppressed individuals (**Fig. 1g, h; Extended Data Fig. 2a**). Moreover, we observed robust proliferative responses to structurally networked epitopes, including a highly proliferative CD8^+^ T cell response to the networked B57:01-restricted TW10 Gag epitope in the San Francisco EEC-I patient and a B*07:02-restricted TL10 Nef response in an EEC-II, whereas the corresponding response in an ART-suppressed individual was markedly lower (**Fig. 1g**). When the analysis was restricted to networked epitopes, both the magnitude and breadth of proliferative responses were significantly elevated in exceptional controllers (**Fig. 1i, j**), illustrating that EECs are distinguished by both the proliferative capacity and specificity of their CD8⁺ T cell responses.

We next evaluated the sequence variation of targeted epitopes within autologous proviral clones. Across all epitopes, per-residue mutational frequencies did not differ significantly between EECs and ART-suppressed individuals (**Fig. 1k**). However, within exceptional controllers, networked epitopes accumulated markedly fewer mutations than non-networked epitopes (**Fig. 1l**), despite robust networked epitope-specific CD8^+^ T cell targeting. In addition, given prior evidence that the integrase K258R mutation can bias HIV integration toward centromeric regions^21^, we evaluated the integrase K258 residue as a potential contributor to the reservoir configuration observed in EECs^6^. Integrase K258 remained wild type in all evaluated EEC (n = 10) and ART (n = 20) proviral sequences (**Fig. 1m**), making K258R-mediated preferential centromeric integration an unlikely explanation for the unique reservoir phenotype. Moreover, in the two EEC-IIs in whom autologous reservoir variants were detected within immunodominant networked epitopes, these variant epitopes elicited CD107a^+^ IFN-γ^+^ CD8^+^ T cell responses comparable to those induced by the corresponding consensus peptides (**Fig. 1n, o**). Collectively, these data demonstrate that EECs preferentially generate proliferative CD8⁺ T cell responses against networked HIV epitopes that are conserved or remain cross-recognizable in autologous virus, establishing an epitope-specific basis for exceptional control.

### An NK-like transcriptional program distinguishes HIV-specific CD8⁺ T cells in EECs

To define the transcriptional states of networked epitope-specific CD8⁺ T cells in exceptional controllers, we performed single-cell RNA sequencing of sorted multimer-positive CD8⁺ T cells specific for networked epitopes from EECs (n = 8) and ART-suppressed individuals (n = 3) (**Fig. 2a; Extended Data Fig. 3**). For EECs, cells were profiled both in the presence and absence of peptide stimulation, enabling comparison of *ex vivo* and expanded antigen-specific populations. Unsupervised clustering identified 17 transcriptionally distinct clusters across networked HIV-specific CD8⁺ T cells (**Fig. 2b**), none of which was biased by a single individual (**Extended Data Fig. 4a**). Analysis of group composition across clusters revealed that clusters 16, 3, and 14 were the most enriched CD8^+^ T cell subsets within both unstimulated and stimulated EEC-derived cells (**Fig. 2c, Extended Data Fig. 4b**). While clusters 3 and 14 exhibited effector and stem-like memory CD8^+^ T cell transcriptional profiles (**Extended Data Fig. 4c, d**) that have previously been implicated in elite and post-treatment control^22–24^, the EEC-enriched cluster 16 was unexpectedly marked by a prominent NK-like transcriptional program, with elevated expression of KIR transcripts, such as *KIR2DL1*, *KIR2DL3*, *KIR3DL1* and *KIR3DL2*, as well as *FCGR3A, TYROBP, IKZF2, CX3CR1*, and *KLRC* family members (*KLRC1, KLRC2,* and *KLRC3*) (**Fig. 2d, e**). The cytotoxic module score, including genes *PRF1, GZMB, NKG7,* and *GNLY*, was also significantly enriched in cluster 16 **(Extended Data Fig. 4e, Extended Table 2)**. Moreover, gene set enrichment analysis (GSEA) revealed that cluster 16 had a transcriptional program enriched for tumor necrosis factor (TNF)-α signaling and interferon (IFN)-α response pathways **(Extended Data Fig. 4f)**, whereas other clusters were enriched for cell cycle pathways. TNF-α, IFN-α, and IFN-γ response pathways were further enriched in cluster 16 cells from EECs compared with ART individuals, supporting enrichment of inflammatory and interferon-responsive programs within this NK-like subset in EECs **(Extended Data Fig. 4g)**. To determine whether the enrichment of this HIV-specific CD8^+^ T cell subset within EECs persisted even after accounting for peptide stimulation, we directly compared stimulated EEC and ART-suppressed transcriptional profiles. This analysis confirmed that the NK-like cluster remained the most strongly enriched population in EECs (**Fig. 2f**). These data indicate that networked HIV-specific CD8⁺ T cell responses in exceptional controllers are distinguished from responses in ART-suppressed individuals not only by their memory and proliferative features, but also by the enrichment of cells that exhibit an NK-like cytotoxic program.

**Figure 2.**
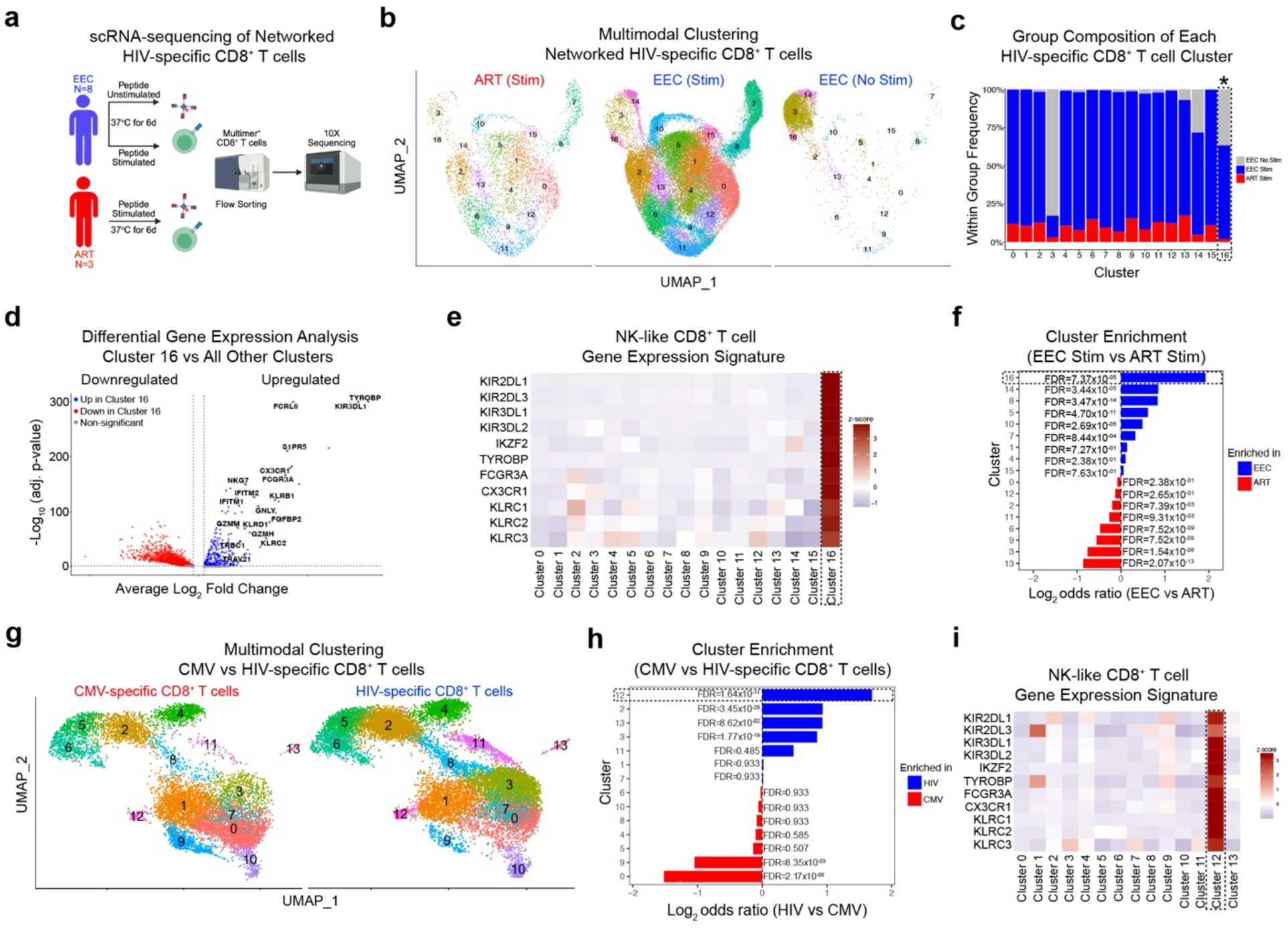
A cytotoxic NK-like CD8⁺ T cell state is selectively enriched within HIV-specific responses in exceptional controllers. **a**, Schematic overview of single-cell RNA sequencing of networked HIV-specific CD8⁺ T cells from EECs (n = 8) and ART individuals (n = 3). PBMCs were stimulated with mapped networked epitopes for 6 days or left unstimulated, and multimer-positive networked HIV-specific CD8⁺ T cells were sorted for 10x Genomics 5′ scRNA-seq. **b,** UMAP of all networked HIV-specific CD8⁺ T cells, colored by cluster identity and separated by condition (ART stimulated, EEC stimulated, and EEC unstimulated). **c,** Frequency of each transcriptional cluster within the total networked HIV-specific CD8⁺ T cell pool for each group. **d,** Volcano plot of differential gene expression comparing cluster 16 with all other clusters. Top differentially expressed genes are labeled with selected NK-lineage genes highlighted. **e,** Heatmap showing average scaled expression of selected NK cell signature-associated genes (*KIR2DL1*, *KIR2DL3*, *KIR3DL1*, *KIR3DL2*, *FCGR3A*, *TYROBP*, *CX3CR1*, *KLRC1*, *KLRC2*, *KLRC3*, *IKZF2*) across all clusters. **f,** Cluster enrichment analysis in stimulated EEC versus ART cells. Bars show log2 odds ratios with significance assessed by Fisher’s exact test with FDR correction. **g,** UMAP of CMV-specific (pp65 NV9) and HIV-specific (Gag SL9 or RT IV9) CD8⁺ T cells sorted from the same EC and EEC-II donors (n = 4; ECs, n = 2; EEC-IIs, n = 2) colored by cluster identity and separated by antigen specificity. **h,** Cluster enrichment analysis comparing HIV-specific versus CMV-specific CD8⁺ T cells. Bars show log₂ odds ratios; significance assessed by Fisher’s exact test with FDR correction. **i,** Heatmap showing average scaled expression of selected NK-lineage genes across CMV-specific and HIV-specific CD8⁺ T cell clusters.

To explore the developmental relationships among NK-like CD8^+^ T cells and other networked HIV-specific CD8^+^ T cell states, we examined T cell receptor (TCR) clonotypes and transcriptional relatedness across clusters. TCR repertoire analysis revealed that the NK-like CD8^+^ T cell cluster had an oligoclonal repertoire, with more than 75% of cells belonging to large or hyperexpanded clones **(Extended Data Fig. 5a)**. Shared clonotype analysis demonstrated significant overlap between the NK-like cluster 16 and memory CD8^+^ T cell cluster 3 **(Extended Data Fig. 5a, b)**, accompanied by partial transcriptional overlap **(Extended Data Fig. 5c)**. Using cluster 3 as the root state for trajectory inference, we found that the NK-like CD8^+^ T cell cluster aligned along a differentiation trajectory from memory HIV-specific CD8^+^ T cells **(Extended Data Fig. 5d)**. In addition, antigen stimulation induced robust expansion of dominant TCR clonotypes across EEC donors **(Extended Data Fig. 5e**). These findings suggest that the NK-like state is embedded within an antigen-experienced, clonally expanded HIV-specific CD8^+^ T cell repertoire.

We next asked whether this NK-like signature was restricted to networked epitope-specific CD8^+^ T cell responses or represented a broader feature of HIV-specific CD8⁺ T cell immunity along the spectrum of elite and exceptional HIV control. To address this, we analyzed an independent cohort of elite and exceptional controllers (n = 4; EC, n = 2; EEC-II, n = 2) in which immunodominant HLA-A*02-restricted HIV-specific (Gag p24 SL9 or RT IV9)- and HLA-A*02-restricted cytomegalovirus (CMV)-specific (pp65 NV9) CD8⁺ T cells were sorted from the same individuals using epitope-specific HLA multimers and profiled by single-cell RNA sequencing (**Extended Data Fig. 3b**). Unsupervised clustering identified 14 transcriptionally distinct clusters (**Fig. 2g**). Direct comparison of these antiviral CD8^+^ T cell specificities revealed preferential enrichment of an NK-like CD8^+^ T cell cluster within HIV-specific CD8⁺ T cells relative to CMV-specific CD8⁺ T cells within the same individuals (**Fig. 2h**). This HIV-enriched cluster expressed a transcriptionally analogous NK-like gene signature, including canonical *KIR* transcripts, *TYROBP*, *FCGR3A*, *CX3CR1*, and *KLRC* family members (**Fig. 2i**). Thus, the NK-like transcriptional program first observed within networked HIV epitope-specific responses extends more broadly across EC and EEC HIV-specific CD8⁺ T cell responses.

### Broader KIR^+^ CD8^+^ T cells in exceptional controllers exhibit an interferon-primed NK-like program

Having identified an NK-like transcriptional program in both networked epitope-specific and immunodominant HLA-A*02-restricted HIV-specific CD8^+^ T cells, we next asked whether EECs harbor a broader KIR^+^ CD8^+^ T cell compartment with related cytotoxic and antiviral transcriptional features. We therefore performed single-cell RNA sequencing of bulk KIR^+^ CD8 T cells sorted from a subgroup of EECs (n = 4) and ART individuals (n = 5) using a cocktail of KIR2DL1, KIR2DL2/L3, KIR3DL1, and KIR3DL2-specific antibodies **(Fig. 3a; Extended Data Fig. 6a)**. Unsupervised clustering identified 13 transcriptionally distinct subsets within the KIR⁺ CD8⁺ T cell compartment **(Fig. 3b)**, all of which expressed KIR transcripts **(Extended Data Fig. 6b).** Among these clusters, cluster 5 was the most strongly enriched in EECs relative to ART-suppressed individuals and expressed canonical NK cell transcripts (*KIR*, *KLRC* genes) and cytotoxic effector genes (*NKG7*, *GNLY, GZMB*, and *PRF1*) **(Fig. 3c, d; Extended Data Fig. 7a)**. To determine whether any KIR^+^ CD8^+^ T cell subset resembled the NK-like HIV-specific CD8^+^ T cell cluster (cluster 16; **Fig. 2**), we calculated a module score based on the top 100 differentially expressed cluster 16 genes **(Extended Table 3)**. This analysis identified EEC-enriched cluster 5 as the KIR^+^ CD8^+^ T cell subset with the highest median enrichment score (0.663; Kruskal-Wallis test, χ^2^ = 40,244, df = 12) **(Fig. 3e)**, indicating transcriptional overlap with HIV-specific NK-like CD8^+^ T cells. In addition, KIR^+^ CD8 T cells from EECs exhibited significantly higher NK cell module scores (calculated using NK cell marker and effector gene sets) **(Fig. 3f)**, whereas KIR^+^ CD8 T cells from ART-suppressed individuals had a memory T cell phenotype **(Fig. 3g)**.

**Figure 3.**
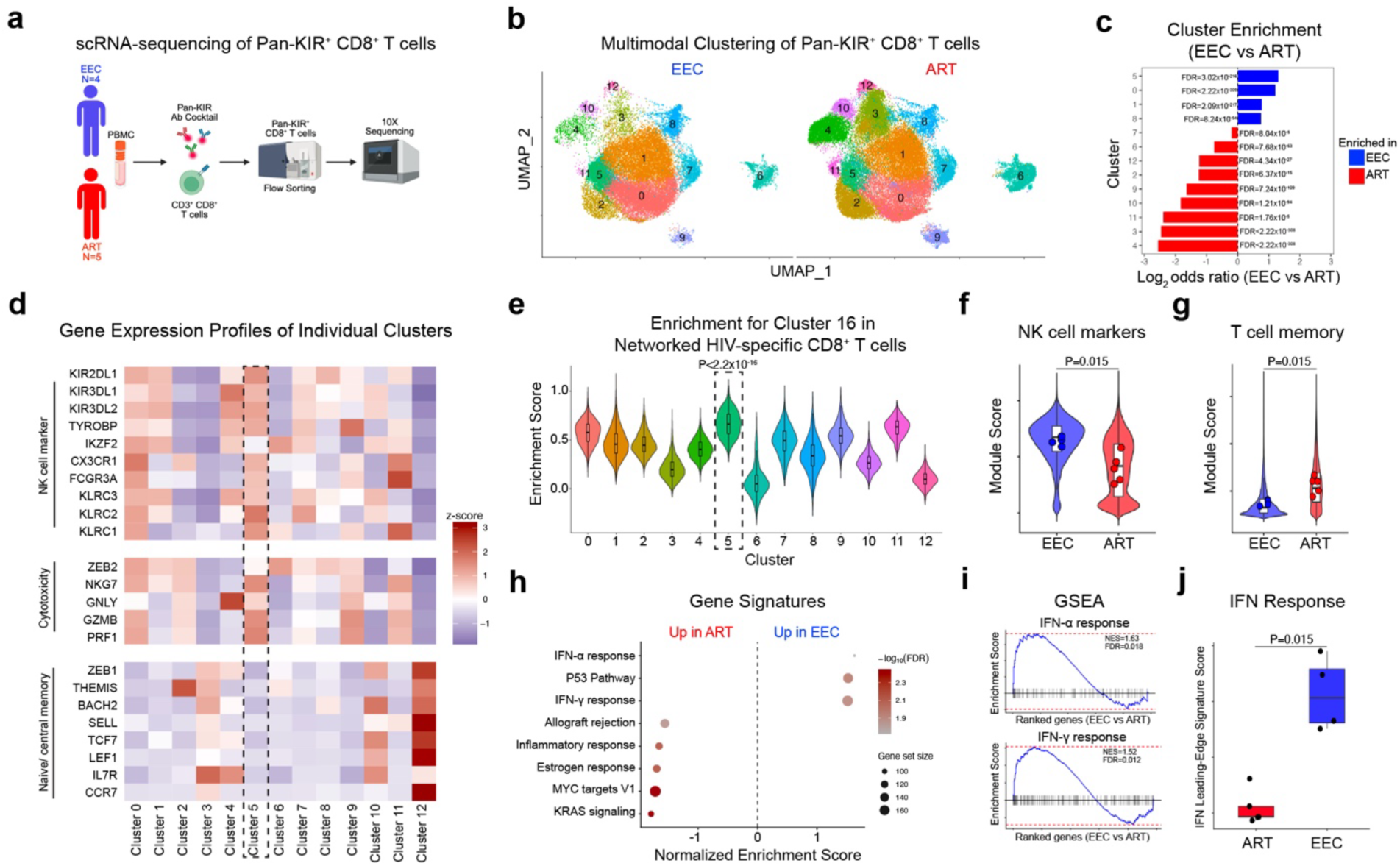
KIR⁺ CD8⁺ T cells from exceptional controllers adopt an interferon-primed NK-like transcriptional profile. **a**, Schematic of single-cell RNA sequencing of total pan-KIR⁺ CD8⁺ T cells sorted from EECs (n = 4) and ART individuals (n = 5) using a cocktail of KIR2DL1, KIR2DL2/L3, KIR3DL1, and KIR3DL2 antibodies. **b,** UMAP of 13 transcriptionally distinct clusters within the global KIR⁺ CD8⁺ T cell compartment, colored by cluster identity. **c,** Cluster enrichment analysis comparing EEC versus ART KIR⁺ CD8⁺ T cells. Bars show log_2_ odds ratios with significance assessed by Fisher’s exact test with FDR correction. **d,** Heatmap showing average scaled expression of canonical NK cell and cytotoxic effector genes (*KIR* family, *KLRC* family, *NKG7*, *GNLY*, *GZMB*, *PRF1*) across all clusters. **e,** Module score comparison of global KIR⁺ CD8⁺ T cell clusters with the NK-like HIV-specific cluster 16 (Fig. 2), calculated using the top 100 differentially expressed genes from cluster 16. **f, g,** NK cell and memory T cell module scores in KIR⁺ CD8⁺ T cells from EECs versus ART individuals. **h,** Normalized enrichment scores for gene signatures within cluster 5 comparing EEC versus ART. **i**, Gene set enrichment analyses for IFN-α and IFN-γ response pathways. **j,** IFN-response leading-edge module score in KIR⁺ CD8⁺ T cells from EECs versus ART individuals, derived from leading-edge genes of significant IFN-α and IFN-γ GSEA pathways.

We next focused on cluster 5 to further define transcriptional differences between EEC- and ART-derived NK-like KIR^+^ CD8^+^ T cells. Differential expression analysis identified increased expression of interferon-inducible genes (*GBP5*)^25^ in EEC cluster 5 cells (**Extended Data Fig. 7b**). Gene set enrichment analyses (GSEA) further demonstrated type I and type II interferon responses in EEC cluster 5 cells, with significant enrichment of the IFN-α and IFN-γ response gene sets **(Fig. 3h, i),** consistent with an antiviral program. In contrast, cells from ART participants were enriched for MYC targets and KRAS signaling, which likely reflects a metabolic remodeling state. To quantify the IFN program driving this enrichment in EECs, we extracted the leading-edge genes from IFN-α and IFN-γ response pathways (**Extended Data Fig. 7c)** and generated an IFN-response module score. Consistent with pathway analyses, the IFN leading-edge module score was significantly higher in cluster 5 cells from EECs than in ART individuals **(Fig. 3j),** paralleling the IFN-responsive features observed in NK-like HIV-specific CD8^+^ T cells. Collectively, these findings indicate that EECs harbor a KIR^+^ CD8⁺ T cell compartment enriched for cytotoxic and IFN-responsive programs transcriptionally related to HIV-specific NK-like CD8^+^ T cells, providing the basis for protein-level and functional assessments of KIR^+^ CD8^+^ T cells in exceptional controllers.

### KIR⁺ CD8⁺ T cells exhibit elevated frequencies in EECs and contribute non-redundantly to the elimination of HIV-infected CD4⁺ T cells

Given the transcriptional evidence for an NK-like cytotoxic program within KIR^+^ CD8^+^ T cells in EECs, we next assessed KIR^+^ CD8^+^ T cell frequencies and their contribution to the elimination of HIV-infected CD4^+^ T cells. Importantly, we gated on CD3^+^ CD8^+^ T cells and excluded unconventional T cells, such as invariant NK T cells, mucosal-associated invariant T (MAIT) cells, and ψο T cells (**Extended Data Fig. 8a**). Representative staining showed higher frequencies of KIR⁺ CD8⁺ T cells in EECs (n = 10) than in ART-suppressed individuals (n = 10) (**Fig. 4a,b**). In light of our scRNA-sequencing data (**Fig. 2, 3**), which demonstrated co-expression of *KIR* transcripts and *KLRC* genes (encoding surface receptors NKG2A and NKG2C), we also evaluated for the expression of NKG2A and NKG2C on CD8^+^ T cells (**Fig. 4a**) in a subset of EECs (n = 8) with sufficient sample availability. Although prior work has suggested that KIR and NKG2A are expressed on distinct, non-overlapping CD8^+^ T cell populations in humans^26^, we surprisingly observed that these EECs were enriched for CD8⁺ T cells expressing both KIR and NKG2A (**Fig. 4a,c**). In contrast, frequencies of dual KIR⁺ NKG2C⁺ CD8⁺ T cells, which can expand in response to human CMV infection^27^, did not differ significantly between groups (**Fig. 4d**). These data demonstrate at a protein level that exceptional controllers are enriched for KIR^+^ CD8⁺ T cells and identify dual KIR and NKG2A expression as a distinguishing feature.

**Figure 4.**
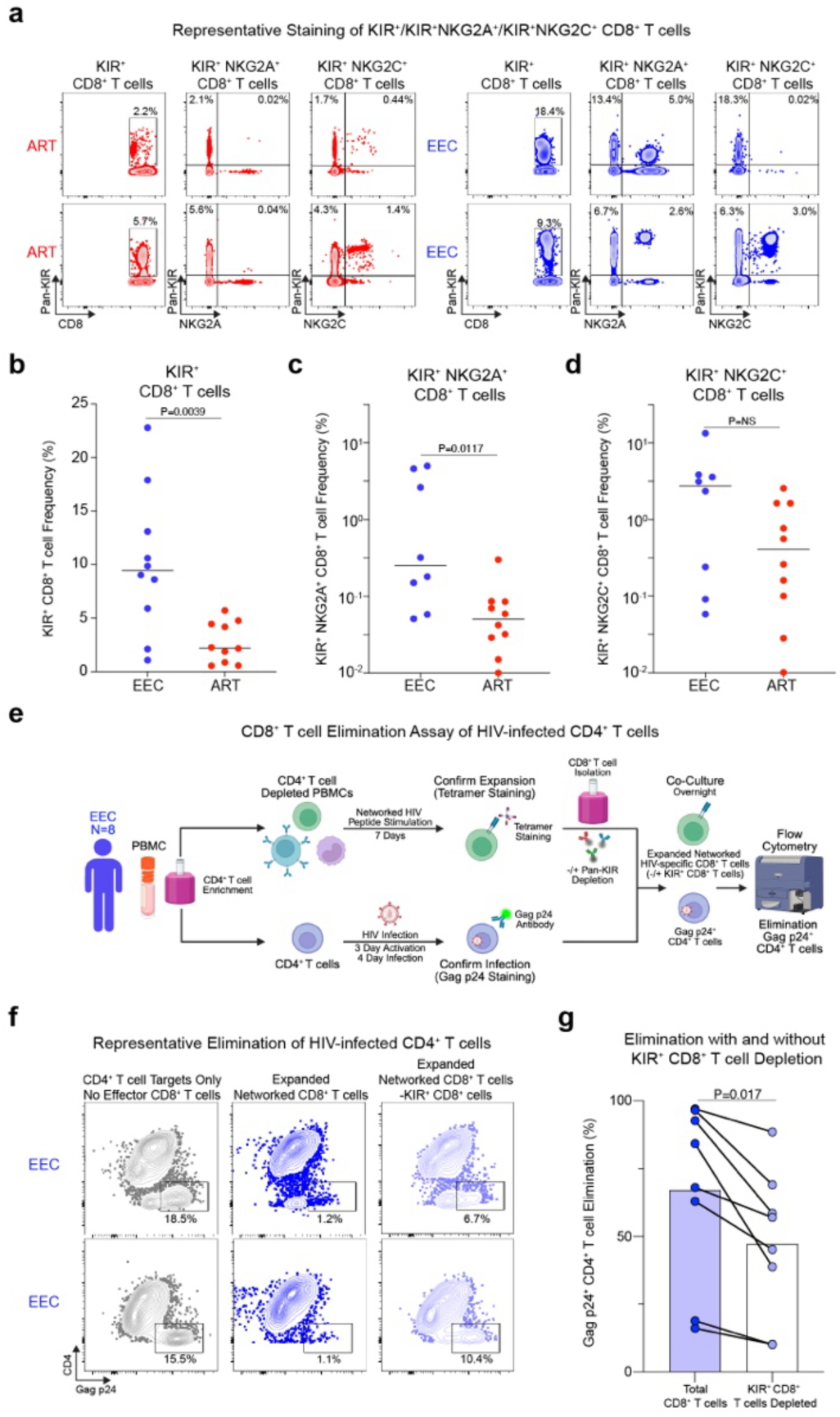
KIR⁺ and KIR⁺NKG2A⁺ CD8⁺ T cells are expanded in exceptional controllers and functionally contribute to autologous HIV-infected cell elimination. **a**, Representative flow cytometry plots showing surface expression of pan-KIR, NKG2A, and NKG2C on CD8⁺ T cells from representative EEC and ART-suppressed individuals. **b,** Comparison of frequencies of pan-KIR⁺ CD8⁺ T cells in EECs (n = 10) versus ART-suppressed individuals (n = 10). **c,** Comparison of frequencies of dual KIR⁺NKG2A⁺ CD8⁺ T cells in EECs (n = 8) versus ART-suppressed individuals (n = 10). **d,** Comparison of frequencies of dual KIR⁺NKG2C⁺ CD8⁺ T cells in EECs (n = 8) versus ART-suppressed individuals (n = 10). **e,** Schematic of the autologous HIV elimination assay. CD4⁺ T cells from EECs were activated and infected with HIV-1 NL4-3. Matched PBMCs were expanded with mapped networked epitope peptides for 6 days, and CD8⁺ T cells were purified by positive selection. KIR⁺ cells were depleted using an APC-conjugated KIR antibody cocktail and anti-APC microbeads. Total CD8⁺ or KIR-depleted CD8⁺ effectors were co-cultured with HIV-infected CD4⁺ target cells at a 1:1 effector-to-target ratio for 3 days, followed by flow cytometric quantification of residual Gag p24⁺ CD4⁺ T cells. **f,** Representative flow cytometry plots showing Gag p24 staining in HIV-infected CD4⁺ target cells co-cultured with no effectors, total CD8⁺ T cells, or KIR-depleted CD8⁺ T cells for two representative EEC donors. **g,** Percentage elimination of autologous HIV-infected CD4⁺ T cells by total versus KIR-depleted CD8⁺ T cells across eight EEC donors. Percent elimination = [1 − (%p24⁺ with effector / %p24⁺ without effector)] × 100. Paired comparisons were made using the Wilcoxon signed-rank test.

To determine whether KIR⁺ CD8⁺ T cells contribute to antiviral function, we developed an autologous elimination assay using HIV-infected CD4⁺ T cells as targets and matched PBMCs stimulated with mapped networked HIV epitopes as antigen-specific CD8⁺ T cell effectors (**Fig. 4e, Extended Data Fig. 8b**). Expanded networked epitope-specific CD8⁺ T cells were then co-cultured with infected CD4^+^ T cells as either total CD8⁺ T cells or after depletion of KIR^+^ CD8^+^ T cells (**Extended Data Fig. 8c-e**). Depletion efficiency and post-sort culture purity were confirmed by flow cytometry, and effector-to-target (E:T) ratios were normalized after depletion. Total expanded CD8⁺ T cells efficiently eliminated autologous Gag p24⁺ CD4⁺ T cell targets in representative EEC donors (1:1 E:T ratio), reducing the frequency of infected cells from 18.5% to 1.2% and from 15.5% to 1.1%, respectively (**Fig. 4f**). In contrast, depletion of KIR⁺ CD8⁺ T cells impaired target cell elimination, with residual Gag p24⁺ CD4⁺ T cells increasing to 6.7% and 10.4% in the same donors despite preservation of the expanded networked epitope-specific effector population. Across eight EEC donors, removal of KIR⁺ CD8⁺ T cells led to a significant reduction in the elimination of autologous HIV-infected CD4⁺ T cells (**Fig. 4g**). This functional deficit demonstrates that KIR⁺CD8⁺ T cells are not merely a phenotypic correlate of exceptional control but contribute non-redundantly to the cytolytic activity of expanded HIV-specific CD8⁺ T cell populations. These findings suggest that NK-like KIR⁺ CD8⁺ T cells serve as a functional effector axis that complements proliferative networked epitope-specific CD8⁺ T cell immunity in exceptional HIV control.

## Discussion

In this study, we characterized the cellular immune features associated with exceptional HIV control. We find that EECs retain canonical features of effective antiviral CD8⁺ T cell immunity, including robust proliferative capacity directed toward structurally constrained networked epitopes^1,2^, which are conserved or remain cross-recognizable in autologous provirus. In addition, we observed that HIV-specific CD8⁺ T cell subsets in EECs are enriched for both effector and stem-like memory CD8^+^ T cell populations, consistent with prior work of elite and post-treatment control^22–24^. However, we also identified the marked enrichment of a unique NK-like CD8^+^ T cell subset, characterized by expression of inhibitory KIR molecules, *KLRC/NKG2*, *IKZF2*, *FCGR3A*, *TYROBP*, and cytotoxic effector genes. This CD8^+^ T cell state was first detected within networked HIV epitope-specific responses but was subsequently observed more broadly in immunodominant HLA-A*02-restricted HIV-specific CD8⁺ T cells and shared transcriptional features with highly cytotoxic subsets within the broader KIR^+^ CD8^+^ T cell compartment, while retaining features of a virus-specific NK-like effector state. Functional depletion experiments demonstrated that KIR⁺ CD8⁺ T cells contribute non-redundantly to the elimination of autologous HIV-infected CD4⁺ T cells. These findings support a model in which exceptional HIV control reflects coordinated activity between proliferative, stem-like memory CD8^+^ T cell responses targeting structurally constrained HIV epitopes and NK-like cytotoxic effector function.

Prior work has shown that CD8^+^ T cells with an NK-like phenotype play roles in transplant rejection^28^ and melanoma clearance^29^, where they utilize rapid and potent cytolytic function. In exceptional controllers, this subset may similarly facilitate efficient elimination of HIV-infected CD4^+^ T cells expressing intact provirus, particularly when paired with sustained recognition of mutationally constrained networked epitopes. Such activity may contribute to immune selection of the reservoir^16^, favoring the unique configurations observed in exceptional controllers^6^, in which intact proviral DNA is confined to transcriptionally repressed genomic regions. Importantly, while this cure-like reservoir state is a defining feature of exceptional controllers, this designation does not imply that they represent a fully discrete immunologic entity from clinically defined elite controllers. Many ECs have not undergone comparable intact proviral sequencing and integration-site analysis and therefore cannot be definitively classified along this reservoir-defined spectrum. Thus, the NK-like CD8^+^ T cell state identified here may reflect a cellular program associated with progressive immune-mediated reservoir restriction across elite control, with reservoir-defined exceptional controllers representing the subset of individuals linked to the most profound restriction or apparent clearance of intact provirus.

The enrichment of interferon-responsive gene elements identified in GSEA of NK-like HIV-specific CD8⁺ T cells and the broader KIR⁺ cytotoxic CD8^+^ T cell population suggests a potential developmental mechanism, consistent with prior work on unconventional CD8^+^ T cells^30^. One possibility is that persistent but controlled antigen exposure may be associated with type I and type II interferon signals that shape both HIV-specific and KIR^+^ CD8⁺ T cells. Accordingly, single-cell trajectory analyses suggest that stem-like HIV-specific CD8⁺ T cells provide a renewable antigen-specific cell subset, while low-level inflammatory cues may promote differentiation into interferon-poised NK-like effectors capable of eliminating HIV-infected target cells. This is consistent with prior work showing that CD45RA⁺ pan-KIR⁺ memory CD8⁺ T cells correlate inversely with HIV DNA during ART and can suppress reactivated HIV through KIR-dependent mechanisms^31^, while other work has implicated KIR3DL1⁺ CD8⁺ T cells in *in vitro* HIV suppression^32^. The emergence of a similar virus-specific CD8⁺ T cell phenotype in individuals who achieve clearance of chronic HBV infection (i.e., functional cure; as described in Lattouf et al.; co-submitted manuscript) suggests that NK-like CD8^+^ T cells may represent a convergent immune solution across persistent viral infections that become fully controlled. Importantly, the cytolytic activity of NK-like KIR^+^ CD8^+^ T cells in exceptional controllers is distinct from recent reports implicating KIR^+^ CD8^+^ T cells as regulatory T cells that suppress autoreactive and activated CD4^+^ T cells in the context of autoimmunity, COVID-19, and pregnancy^33,34^.

These findings may provide further insight into longstanding genetic evidence of inhibitory KIR molecules in HIV control. Protective interactions between inhibitory KIRs and HLA class I ligands, including KIR3DL1 and HLA-Bw4 alleles, have been associated with slower HIV disease progression^20^ and these compound genotypes were enriched within the exceptional controllers in this study. Moreover, polymorphisms within both KIR^35^ and HLA class I alleles, which can affect KIR-HLA interactions^10^, have been linked to differences in HIV control. These associations have primarily been interpreted through NK cells^36,37^, in which KIR molecules can promote functional NK cell licensing and clearance of infected cells^38^. However, inhibitory KIR expression on CD8^+^ T cells has also been suggested to facilitate HIV control by enhancing T cell survival^39^. Our data extend this framework by implicating NK-like KIR⁺ HIV-specific CD8⁺ T cells in the non-redundant elimination of autologous HIV-infected CD4^+^ T cells in EECs. The unexpected enrichment of dual KIR- and NKG2A-expressing CD8^+^ T cells is particularly notable, given prior work suggesting that KIR^+^ CD8^+^ and NKG2A^+^ CD8^+^ T cells are distinct populations in humans^26^. Although both receptors are inhibitory, their co-expression may mark a calibrated cytotoxic CD8⁺ T cell state, in which inhibitory KIR and NKG2A signaling supports enhanced elimination potency while limiting excessive activation.

This study has limitations that warrant discussion. Exceptional controllers are rare, and sample availability limited the number of individuals available for single-cell profiling, autologous viral sequencing, and functional depletion assays. In addition, most analyses were performed using peripheral blood, which may not fully capture immune activity within lymphoid tissues and other anatomic sites of HIV persistence. The elimination assay establishes a non-redundant role for KIR⁺ CD8^+^ T cells within expanded antigen-specific CD8⁺ T cell populations, but the precise mechanism of target cell killing remains to be elucidated. Studies in larger exceptional controller cohorts and perturbations of KIR and NKG2 receptor pathways will be important for defining the effector mechanisms of this NK-like CD8⁺ T cell subset.

In summary, these findings suggest that the cure-like reservoir configuration observed in exceptional controllers may be favored by coordinated induction of proliferative CD8⁺ T cells targeting structurally constrained networked epitopes together with an NK-like KIR⁺ cytotoxic subset capable of enhanced elimination of infected cells. This dual program provides a rational framework for immunotherapeutic strategies aimed at recapitulating durable treatment-free remission in PLWH.

## Methods

### Study participants

This study included exceptional HIV-1 controllers (EEC) with no detectable provirus (type I cure; EEC-I) or proviruses in transcriptionally repressed sites (type II cure; EEC-II)^6^, spontaneous elite controllers (EC), and people living with HIV on antiretroviral therapy (ART) as part of the Ragon HIV positives cohort. Peripheral blood mononuclear cell (PBMC) specimens were obtained from individuals under secondary use study protocols approved by the Massachusetts General Brigham Institutional Review Board (2019P000459, 2014P000661). All human subjects gave written, informed consent. PBMCs from these individuals were collected by Ficoll gradient separation from ACD tubes or leukapheresis samples. They were then cryopreserved and stored in liquid nitrogen for future use. Demographic and clinical characteristics of study participants are reported in Table 1.

### Epitope-specific HIV-specific CD8^+^ T cell IFN-γ ELISpot assay

PBMCs were resuspended at 1 × 10^6^/ml in RPMI-1640 supplemented with 10% FBS (R10) and plated 200 µl per well in Immobilon-P 96-well microtiter plates (Millipore) precoated with 2 µg/ml anti-IFN-γ (clone DK1, Mabtech). Individual HLA-restricted HIV peptides from the optimal A-list^40^ matched to each participant’s HLA genotype were added at 1 µg/ml and incubated at 37°C overnight. Negative control wells did not receive peptide and positive control wells were treated with 1 µg/ml anti-CD3 (clone OKT3, BioLegend) and 1 µg/ml anti-CD28 (clone CD28.8, BioLegend) antibodies. ELISpot assay was performed using manufacturer’s protocol with anti-IFN-γ (clone 1-DK1, Mabtech) capture, biotinylated anti-IFN-γ (clone B6-1, Mabtech) detection, Streptavidin-ALP (Mabtech), and AP Conjugated Substrate (Bio-Rad) followed by disinfection with 0.05% Tween-20 (Thermo Fisher) and automatic analysis using S6 Macro Analyzer (CTL Analyzers). Responses greater than ten spots per well and 3-fold above negative controls were scored as positive.

### Epitope network scores

Epitope network z-scores were calculated as described previously^1^, with weighted sums of residue network scores at HLA anchor and TCR contact amino acid positions within each class I HLA-optimal epitope using structure-based network analysis to predict mutational tolerance from HIV protein crystal structures. Epitopes with network z-scores above 3.06, representing the top quintile, were considered networked.

### Epitope-specific HIV-specific CD8^+^ T cell proliferation assay

This assay was carried out as previously described^1^. Briefly, PBMCs were stained at 37 °C for 20 min with 0.5 µM CellTrace CFSE (Thermo Fisher) as per the manufacturer’s protocol at 1 × 10^6^ cells/ml. Staining was quenched with FBS (Sigma), and cells were washed twice with R10, resuspended at 1 × 10^6^/ml and plated at 200 µl per well in 96-well round-bottom polystyrene plates (Corning). Individual HIV peptides corresponding to IFN-γ ELISPOT responses for each participant were added at 1 µg/ml in duplicate and incubated at 37 °C for 6 days before flow cytometric assessment. Negative control wells received dimethyl sulfoxide (in amounts equivalent to peptide-containing wells), but no peptide. For each assay, three replicate negative control wells were included to ensure a robust assessment of background proliferation. Positive control wells received 1 µg/ml of anti-CD3 (clone OKT3, BioLegend) and anti-CD28 (clone CD28.8, BioLegend) antibodies. On day 6, cells were stained with Live/Dead Violet (Thermo Fisher), PE/Cy7-anti-human CD3 (clone SK7, BioLegend) and APC-anti-human CD8 (clone SK1, BioLegend) and then analyzed by flow cytometry (fig. S4). The frequency of proliferating CD8^+^ T cells was determined by subtracting the average percentage of CD8^+^ T cells in the CFSE_low_ gate following HIV peptide stimulation (for each pair of duplicates) by the maximum percentage of CFSE low cells among the DMSO negative controls.

### Proviral sequencing

Full-length integrated proviral sequencing and quantification of total and intact reservoir sizes were performed as described previously^6,41^. Genomic DNA was extracted from PBMC and diluted to single-genome levels as measured by droplet digital PCR (ddPCR). Diluted DNA was subjected to HIV-1 near full-genome amplification via nested PCR. Near-full-length sequences (>8,000 bp) were sequenced via Illumina MiSeq. Reads were de novo assembled and aligned to HXB2 to identify mutations, such as 5’-defects, large deletions, hypermutation, premature stop codons and internal inversions. Viral sequences lacking such mutations were considered genome intact.

### Variant recognition assay

PBMCs were stimulated for 4 hours with 1 µM of HLA-optimal peptides of clade B HIV consensus and autologous sequences observed in proviral DNA. BV711-conjugated anti-CD107A (clone H4A3, Biolegend) was included during stimulation to measure degranulation. GolgiStop and GolgiPlug (BD Biosciences) were added 2 hours post-stimulation to enable intracellular cytokine staining. Cells were stained with Live/Dead Violet, BV605-conjugated anti-CD3 (clone SK7, Biolegend) and BUV395-conjugated anti-CD8 (clone RPA-T8, BD Biosciences), fixed and permeabilized using Cytofix/Cytoperm (BD Biosciences), stained for intracellular PE-Cy7-conjugated anti-IFN-γ (clone B27, Biolegend) and analyzed via flow cytometry.

### Cell sorting for scRNAseq analysis

PBMCs from each donor were stained for FACS sorting before sequencing. For antigen-specific CD8^+^ T cells, PBMCs were first incubated with Human TruStain FcX blocking reagent (BioLegend, 422302) for 10 min at 4 °C, washed, and incubated with MHC-I dextramers (**Extended Table 4**) for 30 min at room temperature. Cells were then washed and stained with a concentrated antibody mix containing TotalSeq antibodies (**Extended Table 4**) for 30 min at 4 °C. For KIR^+^ CD8^+^ T cells, PBMCs were incubated directly with an antibody mix containing APC-conjugated KIR antibodies (**Extended Table 4**) for 30 min at 4 °C. Cells were then washed twice and stained with 7-AAD viability dye (5 μL; 0.25 μg; Thermo Fisher Scientific, Cat# 00-6993-50) for 5-10 min at room temperature. Without further washing, cells were resuspended in PBS containing 1% BSA and sorted on a BD Aria cell sorter at 1000 cells/μL. For antigen-specific CD8^+^ T cells, cells from 2-10 donors were pooled, yielding ten total pooled sample sets. Sorted cells were resuspended at the concentration recommended by 10x Genomics for targeted cell recovery prior to loading.

### scRNA-seq data processing and analysis

scRNA-seq libraries for antigen-specific CD8^+^ T cells were generated using the 10x Genomics Chromium 5’ workflow and sequenced on an Illumina NovaSeq platform. Libraries for KIR^+^ CD8^+^ T cells were generated using the 10x Genomics Chromium 3’ workflow and sequenced on an Illumina NovaSeq platform. Raw sequencing data were processed using the 10x Genomics Cell Ranger Multi pipeline (v8.0.1) aligned to the GRCh38 reference genome. Downstream analyses were performed in R (version 4.5.1) using Seurat (version 5.4).

Cells with < 200 detected genes were removed. Upper thresholds for detected genes and mitochondrial RNA content were determined based on inspection of quality metric distributions to remove outlier cells and likely doublets and were applied as follows: non-networked antigen-specific CD8^+^ T cells (> 3000 features or > 10% mitochondrial reads), networked antigen-specific CD8^+^ T cells (> 6000 features or > 30% mitochondrial reads), and KIR^+^ CD8^+^ T cells (> 6000 features or > 11% mitochondrial reads). For networked and non-networked antigen-specific CD8^+^ T cells, standard Seurat preprocessing was performed using NormalizeData, FindVariableFeatures, ScaleData (regressing mitochondrial percentage) and RunPCA. For KIR^+^ CD8^+^ T cells normalization was performed using SCTransform (method = glmGamPoi) with regression of mitochondrial and ribosomal gene percentages, followed by RunPCA.

Batch correction across donors was performed using Harmony (version 1.2.4). Corrected embeddings were used for graph construction (FindNeighbors), clustering (FindClusters), and visualization (RunUMAP). Doublets were identified using DoubletFinder (version 2.0.6) with a 5% expected doublet rate. Doublets were removed prior to reclustering. Final clustering resolutions were 1.1 for networked antigen-specific CD8^+^ T cells, 1.0 for non-networked antigen-specific CD8^+^T cells, and 0.5 for KIR^+^ CD8^+^ T cells. For TCR analysis, scRepertoire (v2.4.0) was used, and pseudotime analysis was performed using Slingshot (v2.16.0).

### Cluster enrichment analysis

Cluster enrichment was assessed using Fisher’s exact tests on cluster-centric contingency tables. P values were adjusted using Benjamini-Hochberg method. Directionality of enrichment was summarized using log_2_ odds ratios with Haldane-Anscombe correction.

### Differential expression and module scoring

Cluster-level differential expression was performed using FindAllMarkers (Seurat) with a log_2_ fold-change threshold of 0.25 and a minimum expression fraction (min.pct) of 0.1 (networked cluster 16) and 0.25 (KIR^+^ cluster 5). Heatmaps of average cluster expression were generated using AverageExpression (Seurat) and ComplexHeatmap (version 2.24.1). Gene module scores were calculated using the AddModuleScore function (Seurat).

### Pseudobulk Differential Expression

For EEC versus ART comparisons, pseudobulk RNA-seq profiles were generated by aggregating raw RNA counts per donor using Seurat, treating donors as the unit of replication. The resulting gene-by-sample count matrix was analyzed using DESeq2 (version 1.48.2). Genes with total counts <10 were excluded prior to analysis. Differential gene expression was tested using a negative binomial model with Benjamini-Hochberg correction. Genes with adjusted P <0.05 and |log_2_ fold-change| ≥1 were considered significant.

### Gene Set Enrichment Analysis (GSEA)

Preranked gene set enrichment analysis (GSEA) was performed using fgsea (version 1.34.2). For scRNA-seq analyses, genes were ranked by log_2_ fold-change, whereas pseudobulk analyses used the DESeq2 Wald test statistic. Hallmark gene sets from MSigDB were tested using 10,000 permutations (minSize=15, maxSize=500), and P values were adjusted using the Benjamini-Hochberg correction method. Pathways with adjusted P < 0.05 were considered significant. To define the IFN response signature, leading-edge genes from significant Hallmark interferon pathways were combined. Variance-stabilized pseudobulk expression values (DESeq2 VST) were z-scored per gene, and donor-level signature scores were calculated as the mean expression across genes.

### Assessment of KIR/NKG2A/NKG2C expression on CD8^+^ T cells by flow cytometry

Purified cryopreserved PBMCs were thawed and rested overnight at 37 °C in RPMI-1640 supplemented with 10% FBS, 100 U/mL penicillin, 100 μg/mL streptomycin, and 2 mM L-glutamine (R10 media). Cells were then washed, resuspended in PBS, and stained with live-dead viability dye using LIVE/DEAD™ Fixable Blue Dead Cell Stain (1:1000, Invitrogen Cat# L34962). Cells were then washed and surface-stained at 4 °C for 30 min with defined antibody combinations **(Extended Table 5).** Cells were then fixed and resuspended in PBS. Samples were acquired on a BD Fortessa X20 using BD FACSDiva8.0 (BD Biosciences), and subsequent data analysis was performed using FlowJo 10 (TreeStar). Fluorescence-minus-one (FMO) controls were run with each experiment to determine gates.

### Preparation of HIV-infected CD4^+^ T cell targets

CD4^+^ T cells were enriched from PBMCs using the EasySep Human CD4^+^ T cell Isolation Kit (STEMCELL Technologies, Cat#17952) as per the manufacturer’s instructions. Isolated CD4^+^ T cells were then counted and resuspended at 0.8-1 × 10^6^ cells/mL in R10 containing IL-2 (50 ng/mL). Cells were plated in a 6-well plate and activated with 1μg/ml anti-CD3 (clone OKT3, BioLegend) and 1 μg/ml anti-CD28 (clone CD28.8, BioLegend) antibodies. Activated CD4^+^ T cells were incubated for 4 days at 37 °C. After activation, CD4^+^ T cells were harvested and counted, washed with R10 media, and resuspended at 4 × 10^6^ cells/mL in R10 containing IL-2 (100 U/mL) and polybrene (8 µg/mL, EMD Millipore Cat#: TR-1003-G). Cells were then infected with HIV-1 NL4-3 by spinoculation (1,200 × g, 90 min, 37 °C). Following centrifugation, cells were incubated for 2 h at 37 °C, washed, counted, and adjusted to 0.8-1 × 10^6^ cells/mL in R10 + IL-2 (100 U/mL). Cells were plated into 6-well plates and cultured for 3 days at 37 °C. Uninfected CD4 T cells were used as a control.

### Preparation of effector CD8^+^ T cells

PBMCs were thawed and stimulated with individual HIV peptides (corresponding to IFN-γ ELISPOT mapping) for 6 days. After 6 days of expansion, CD8^+^ T cells were isolated by positive selection using the REAlease CD8 MicroBead Kit following the manufacturer’s instructions (REAlease CD8 MicroBead Kit; Miltenyi, Cat# 130-117-036). Purified CD8^+^ T cells were washed twice in PBS and stained for 30 min at 4°C with an APC-conjugated KIR antibody cocktail (all 1:50): KIR2DL1/KIR2DS5 (clone 143211, R&D Systems Cat# FAB1844A-100; RRID: AB_416855), KIR3DL2/CD158K (clone 539304, R&D Systems Cat# FAB2878A-100; RRID: AB_3648841), KIR3DL1/CD158e (clone DX9, Miltenyi Biotec Cat# 130-092-474; RRID: AB_871612), CD158b1/b2 (clone GL183, Beckman Coulter Cat# A22333; RRID: AB_3662729). Cells were washed twice, and KIR⁺ cells were subsequently depleted using anti-APC MicroBeads according to the manufacturer’s protocol. KIR⁻ CD8^+^ and total CD8^+^ T cells were counted and resuspended at 1×10⁶ cells/mL and used as effector cells.

### HIV-infected cell elimination assay

HIV-infected CD4^+^ T cell targets were counted and resuspended at 1×10⁶ cells/mL and were co-cultured with KIR-depleted or total CD8^+^ T cell effectors at a 1:1 effector: target ratio for 3 days. After incubation, co-cultures were stained in PBS + 2% FBS (50 µL/well) with LIVE/DEAD Fixable Blue Dead Cell Stain (1:1000, Invitrogen Cat# L34962), Pacific Blue CD3 (UCHT1, 1:100, BioLegend, Cat# 300431, RRID: AB_1595437), PE-Cy7CD4 (RPA-T4, 1:100, BioLegend, Cat# 300512, RRID: AB_314080), and BV605 CD8 (SK1, 1:100, BioLegend, Cat# 344742, RRID: AB_2566513) for 30 min at 4°C, then washed twice. Cells were fixed in FOXP3 fixation/perm buffer for ≥1h at 4°C (or overnight), washed twice with perm wash buffer, and stained intracellularly with FITC anti–HIV p24 (KC57, 1:50, Beckman Coulter, Cat# 6604665, RRID: AB_1575989) for 1hr at 4°C. Samples were washed, resuspended, and acquired on a BD Fortessa X20 using BD FACSDiva8.0 (BD Biosciences). Target elimination was quantified by flow cytometry as the percentage of residual p24^+^ target cells. Percent elimination was calculated as: %Elimination = [1 − (%p24^+^ with effector / %p24^+^ without effector)] x 100.

## Statistical Analysis

Prism 10 (GraphPad Software, Version 10.1.1) was used for statistical analysis as follows: the Mann-Whitney U test was used for single comparisons of independent groups, the Wilcoxon signed-rank test was used to compare two paired groups. For multiple groups, statistical significance was assessed using a Kruskal-Wallis test with Dunn’s post hoc correction. The statistical significances are indicated in the figures (*p < 0.05, **p < 0.01, ***p < 0.001, and ****p < 0.0001), and all tests were two-tailed.

## Acknowledgements

We are grateful to the study participants who made this study possible, and the MGH and Ragon Institute clinical staff for participant recruitment and sample collection.

## Author Contributions

GDG conceived and designed the study. AA, CRC, DRC, SB, CK, UA, XL, AV, MAG, and GDG performed experiments. AA, AKT, CRC, DRC, SB, HH, CK, XL, AV, and GDG analyzed data. DRC, ML, XY, BDW, GML, and GDG provided supervision. ML, XY, BDW, and GDG provided funding for this work. AA, AKT, and GDG wrote the initial draft in consultation with GML. All authors contributed to the final draft.

## Extended Data

**Extended Table 1. Individual clinical, HLA and KIR characteristics**

**Extended Table 2. Gene sets used to compute module scores**

**Extended Table 3. Top 100 genes from cluster 16 used to compute enrichment score**

**Extended Table 4. Antibodies used for scRNAseq analysis**

**Extended Table 5. Antibodies used in KIR^+^ CD8^+^ T cell immunophenotyping**

**Extended Data Figs. 1-8**

**Extended Data Figure 1.**
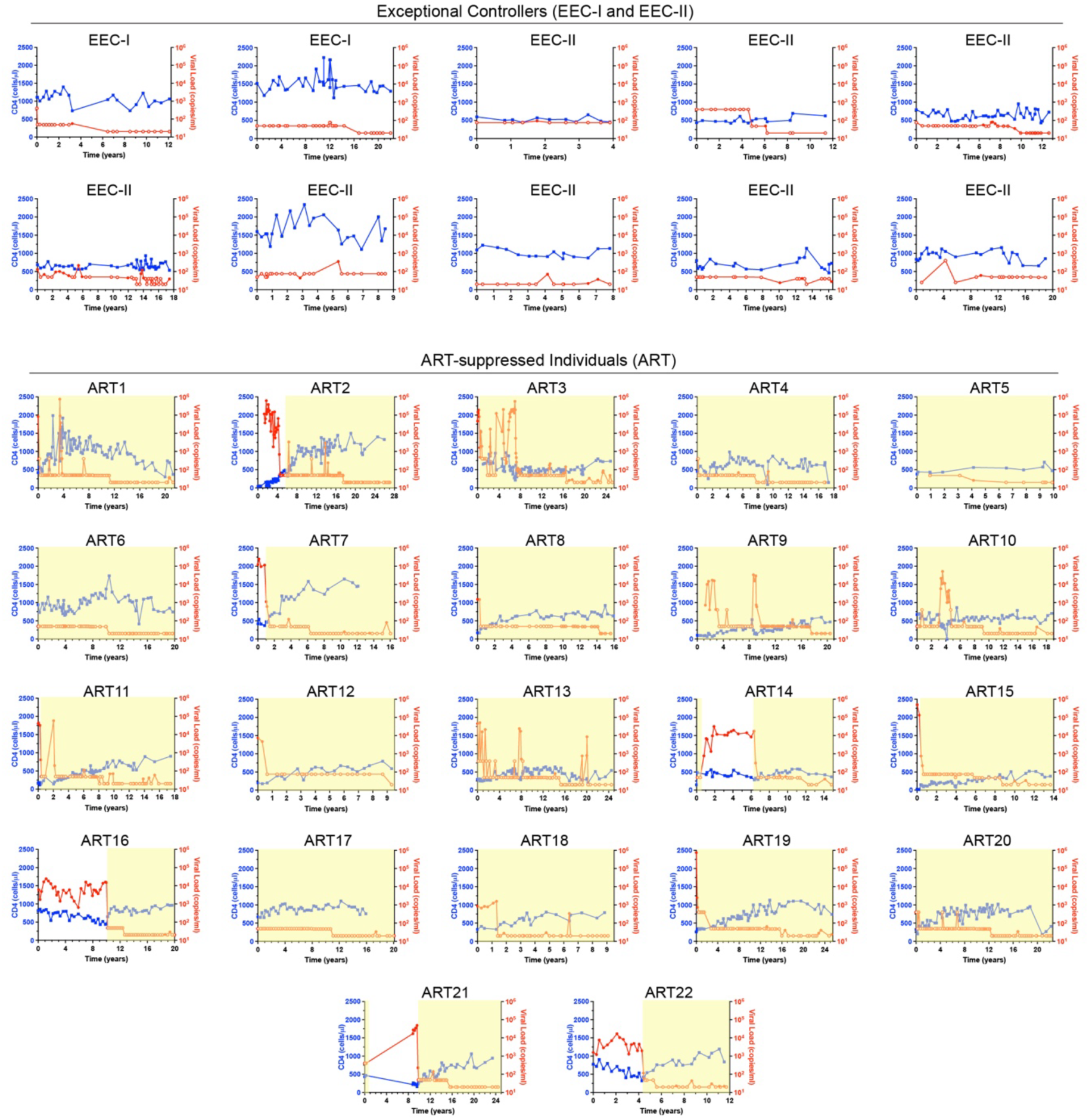
Longitudinal viral load and CD4⁺ T cell trajectories for individual exceptional controllers and ART-suppressed study participants. Individual participant plots showing longitudinal CD4⁺ T cell counts (cells per μl; blue) and viral load (copies per ml; red) over time (years) for all exceptional elite controllers (EEC-I, n = 2; EEC-II, n = 8) and all ART-suppressed individuals (ART; lower panel, n = 22) included in the study. Each panel corresponds to one participant. EEC trajectories demonstrate sustained viral suppression and preserved CD4⁺ T cell counts consistent with exceptional control. ART participant trajectories (highlighted in orange/yellow) show viremic rebound or suppression on therapy. Time zero is defined as the time point of first available clinical data.

**Extended Data Figure 2.**
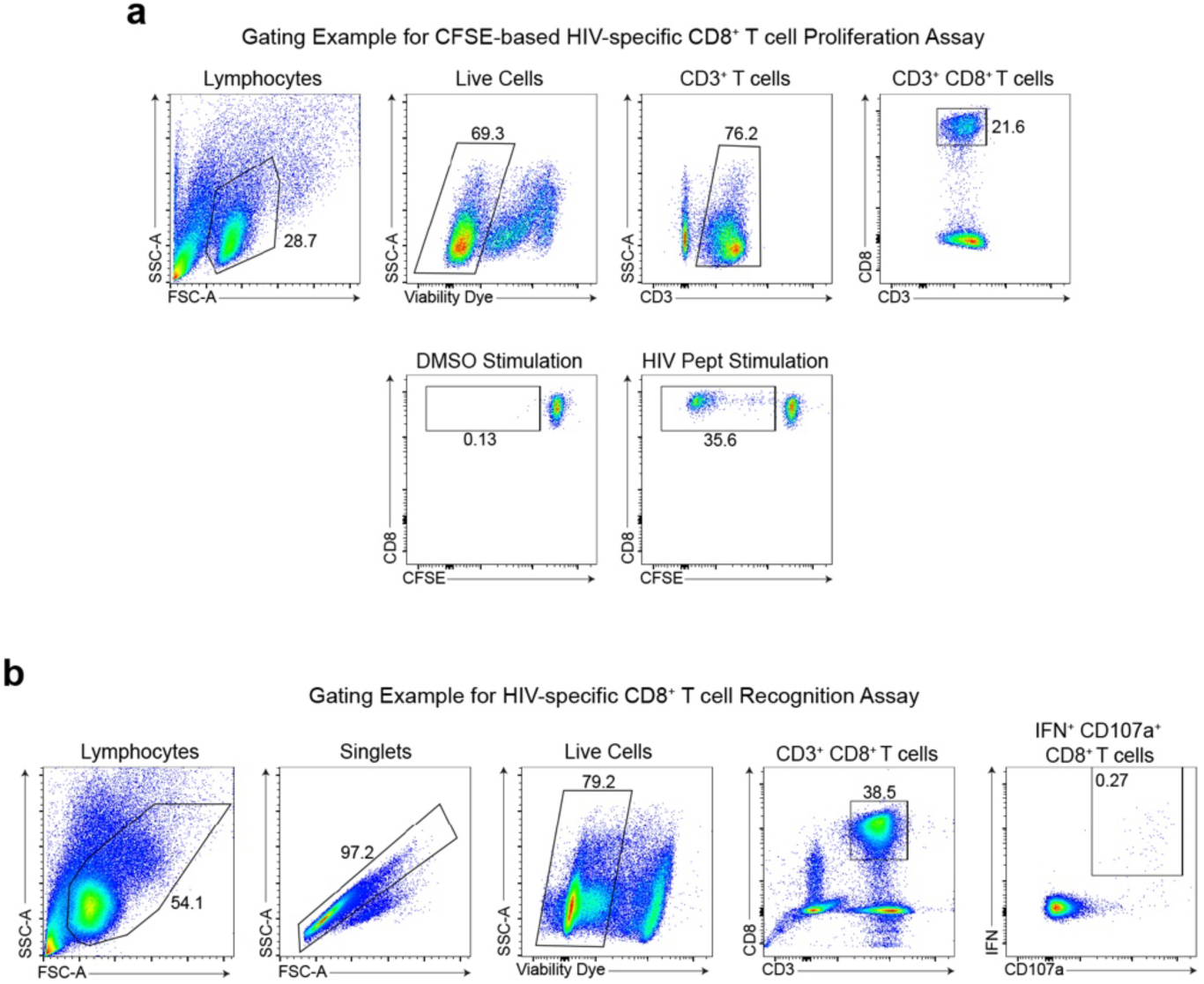
Flow cytometry gating strategies for HIV-specific CD8⁺ T cell proliferation and recognition assays. **a**, Representative gating strategy for the CFSE-based HIV-specific CD8⁺ T cell proliferation assay. Sequential gates are applied to identify lymphocytes (FSC-A vs. SSC-A), live cells (SSC-A vs. viability dye), CD3⁺ T cells (SSC-A vs. CD3), and CD3⁺CD8⁺ T cells (CD8 vs. CD3). CFSE dilution in the CD8⁺ T cell gate is shown for DMSO negative control (0.13% CFSEᴸᴼᵂ) and HIV peptide-stimulated conditions (35.6% CFSEᴸᴼᵂ), illustrating the proliferative CD8⁺ T cell population. **b,** Representative gating strategy for the HIV-specific CD8⁺ T cell recognition assay (IFN-γ/CD107a co-expression). Sequential gates are applied to identify lymphocytes, singlets, live CD3⁺CD8⁺ T cells. HIV-specific CD8⁺ T cells are identified as dual IFN-γ⁺CD107a⁺ in the CD8⁺ T cell gate (representative unstimulated value: 0.27%).

**Extended Data Figure 3.**
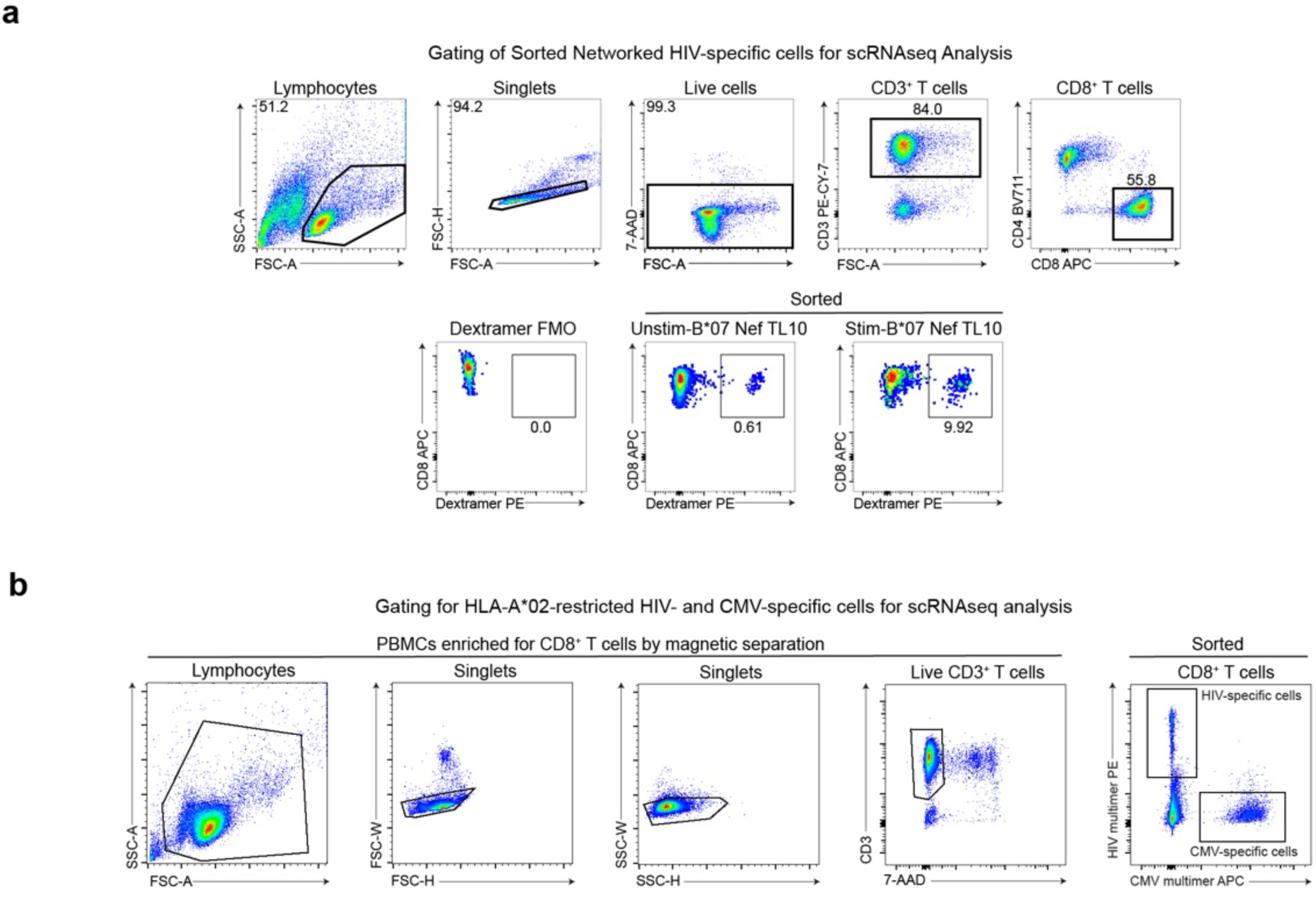
Flow cytometry sorting strategies for single-cell RNA sequencing. **a**, Representative gating strategy for sorting networked HIV-specific CD8⁺ T cells for the primary scRNAseq dataset. Sequential gates identify lymphocytes, singlets, live cells (7-AAD⁻), CD3⁺ T cells, and CD8⁺ T cells. Multimer^+^ networked HIV-specific CD8⁺ T cells are identified using MHC-I dextramers (PE), with a fluorescence-minus-one (FMO) control confirming gate specificity (0.0% in FMO). Representative plots show 0.61% multimer⁺ in unstimulated and 9.92% multimer⁺ in stimulated conditions for a B*07 Nef TL10-specific response. **b,** Representative gating strategy for sorting HLA-A*02-restricted HIV-specific and CMV-specific CD8⁺ T cells for the independent comparative scRNAseq cohort. PBMCs were first enriched for CD8⁺ T cells by magnetic separation. Sequential gates identify lymphocytes, singlets (FSC-H vs. FSC-W and SSC-W vs. SSC-H), live CD3⁺ T cells (7-AAD). HIV-specific (PE-labeled HIV multimer⁺) and CMV-specific (APC-labeled CMV multimer⁺) CD8⁺ T cells are co-sorted into separate populations in a single sort using orthogonal fluorochrome channels.

**Extended Data Figure 4.**
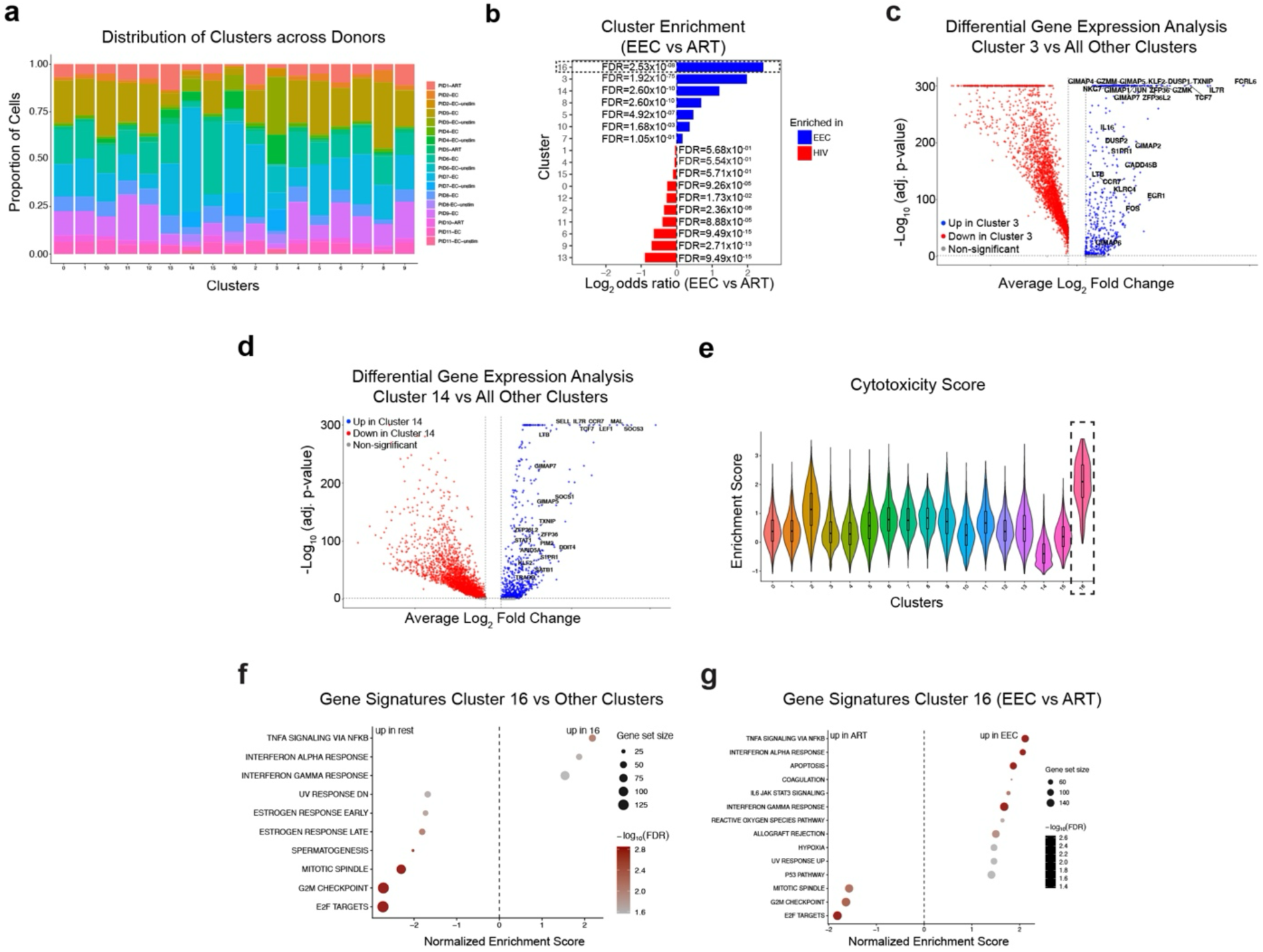
Transcriptional characterization of networked HIV-specific CD8⁺ T cell clusters. **a**, Stacked bar plots showing the proportion of cells from each donor (EEC unstimulated, EEC stimulated, and ART stimulated) within each of the 17 transcriptional clusters, confirming that no single cluster is dominated by cells from a single individual. **b,** Cluster enrichment analysis (log₂ odds ratio, EEC versus ART, stimulated cells) showing the relative enrichment of each cluster in EEC or ART conditions. FDR-adjusted *P* values are indicated for significantly enriched clusters. **c,** Volcano plot of differential gene expression analysis comparing cluster 3 with all other clusters, identifying memory and effector T cell markers (up in cluster 3, shown in blue). **d,** Volcano plot of differential gene expression analysis comparing cluster 14 with all other clusters, identifying canonical stem-like memory T cell markers (up in cluster 14, shown in blue). **e,** Cytotoxicity module score (*PRF1*, *GZMB*, *NKG7*, *GNLY*) across all 17 clusters shown as a violin plot, demonstrating significant enrichment of cytotoxic gene expression in cluster 16 relative to all other clusters (Kruskal-Wallis test, χ² = 8571.8, df = 16, p < 2.2 x 10^-16^). Cluster 16 displayed the highest cytotoxicity score (median = 2.10), significantly exceeding all other clusters (pairwise Wilcoxon tests with Benjamini-Hochberg correction, all adjusted p < 2 x 10^-16^), consistent with a highly cytotoxic CD8^+^ T cells population. **f,** Gene set enrichment analysis (GSEA) bubble plot for cluster 16 versus all other clusters showing top enriched and depleted Hallmark pathways (normalized enrichment scores); IFN-α response and TNF-α signaling pathways are among the top enriched pathways in cluster 16, while cell cycle pathways (mitotic spindle, G2M checkpoint, E2F targets) are depleted. **g,** GSEA bubble plot for cluster 16 cells from EEC versus ART (pseudobulk), showing further enrichment of IFN-α and IFN-γ response gene sets in EEC-derived cluster 16 cells relative to ART-derived cluster 16 cells.

**Extended Data Figure 5.**
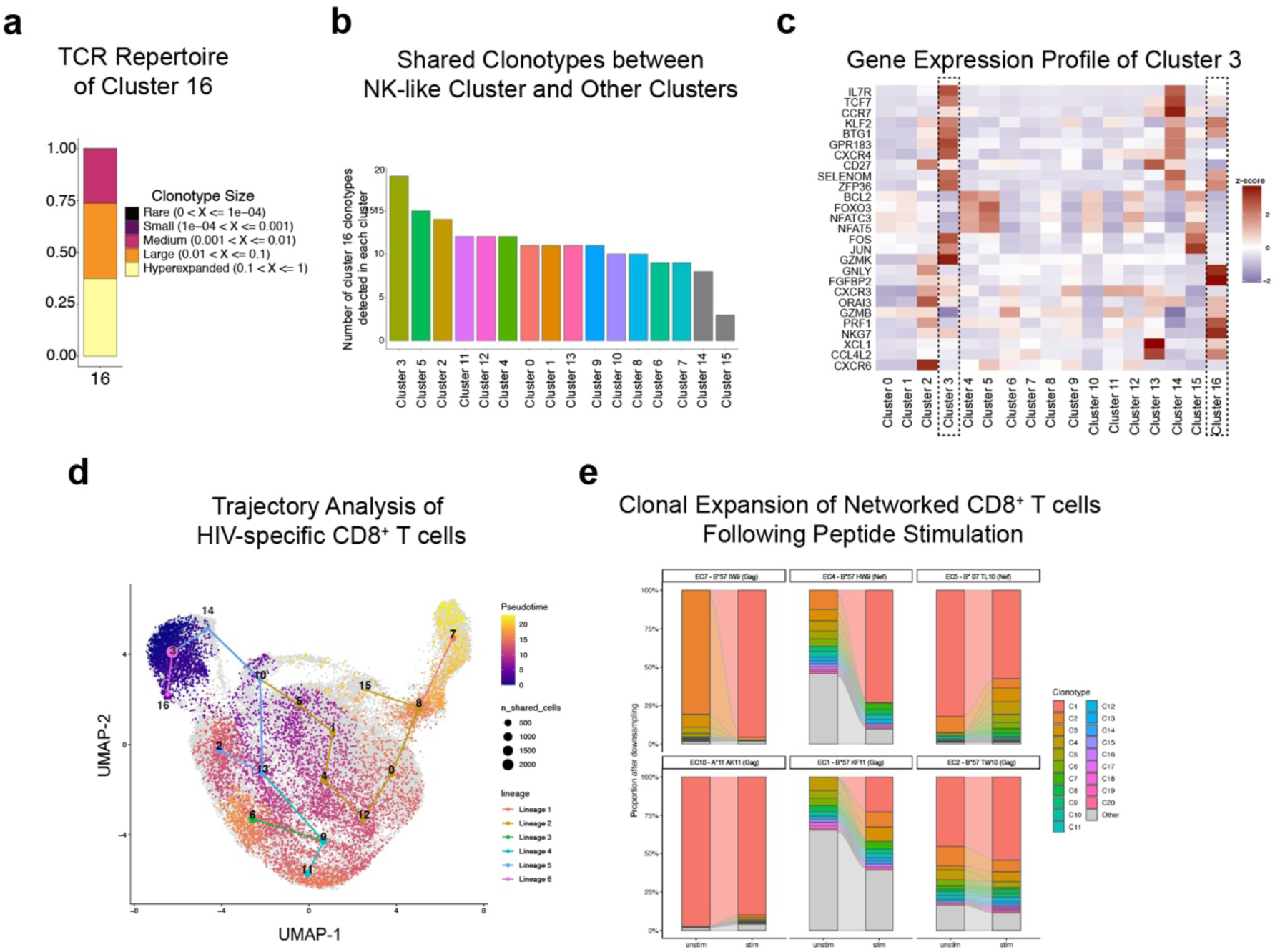
TCR clonotype analysis, differentiation trajectory, and antigen-driven clonal expansion of networked HIV-specific CD8⁺ T cells. **a**, TCR clonotype size distribution within cluster 16 (NK-like), showing that more than 75% of cells belong to large (>10 cells) or hyperexpanded (>100 cells) clones, consistent with an oligoclonal TCR repertoire. **b,** Bar graph showing the number of cluster 16 clonotypes shared with each of the other 16 networked HIV-specific CD8⁺ T cell clusters, demonstrating the greatest clonotype overlap between cluster 16 and cluster 3 (stem-like). **c,** Heatmap of average scaled gene expression for selected transcripts across all clusters, highlighting the stem-like gene expression profile of cluster 3 (high *IL7R*, *TCF7*, *CCR7*, *SELL*, *CD27*, *BTG1*) and the partial transcriptional overlap with NK-like cluster 16. **d,** Slingshot pseudotime trajectory analysis on shared clonotypes between cluster 16 and cluster 3 overlaid on the UMAP of all networked HIV-specific CD8⁺ T cells (gray). Multiple lineage trajectories are shown, with cluster 3 used as the root state. The NK-like cluster 16 differentiates along a direct trajectory from cluster 3, suggesting a stem-to-NK-like differentiation axis. Node size reflects the number of shared clonotypes connecting clusters. **e,** Stacked bar plots showing TCR clonotype composition in unstimulated and antigen-stimulated conditions for six representative EEC donors. Antigen stimulation induces robust expansion of dominant clonotypes (colored bars) relative to unstimulated conditions, confirming antigen-driven clonal proliferation within networked HIV-specific CD8⁺ T cell populations.

**Extended Data Figure 6.**
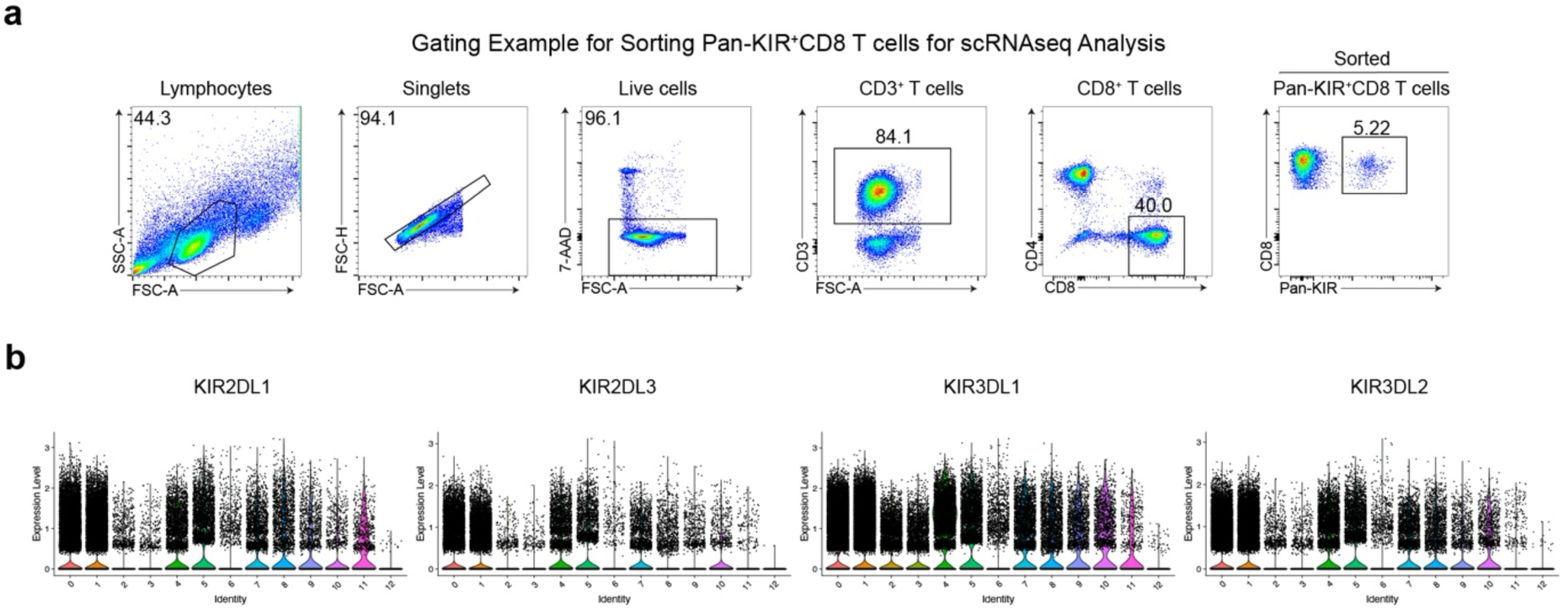
Flow cytometry sorting strategy and KIR transcript expression across global KIR⁺ CD8⁺ T cell clusters. **a**, Representative gating strategy for sorting pan-KIR⁺ CD8⁺ T cells for global KIR⁺ CD8⁺ T cell scRNAseq. Sequential gates identify lymphocytes (FSC-A vs. SSC-A), singlets (FSC-H vs. FSC-A), live cells (7-AAD⁻), CD3⁺ T cells, CD8⁺ T cells (CD8 vs. CD4), and pan-KIR⁺ CD8⁺ T cells using the APC-conjugated KIR antibody cocktail (5.22% of CD8⁺ T cells in a representative donor). **b,** Violin plots showing expression levels of individual KIR transcript family members (*KIR2DL1*, *KIR2DL3*, *KIR3DL1*, *KIR3DL2*) across all 13 transcriptional clusters within the global KIR⁺ CD8⁺ T cell UMAP, confirming that each cluster individually expresses KIR transcripts. Clusters are colored by identity as shown in Fig. 3b.

**Extended Data Figure 7.**
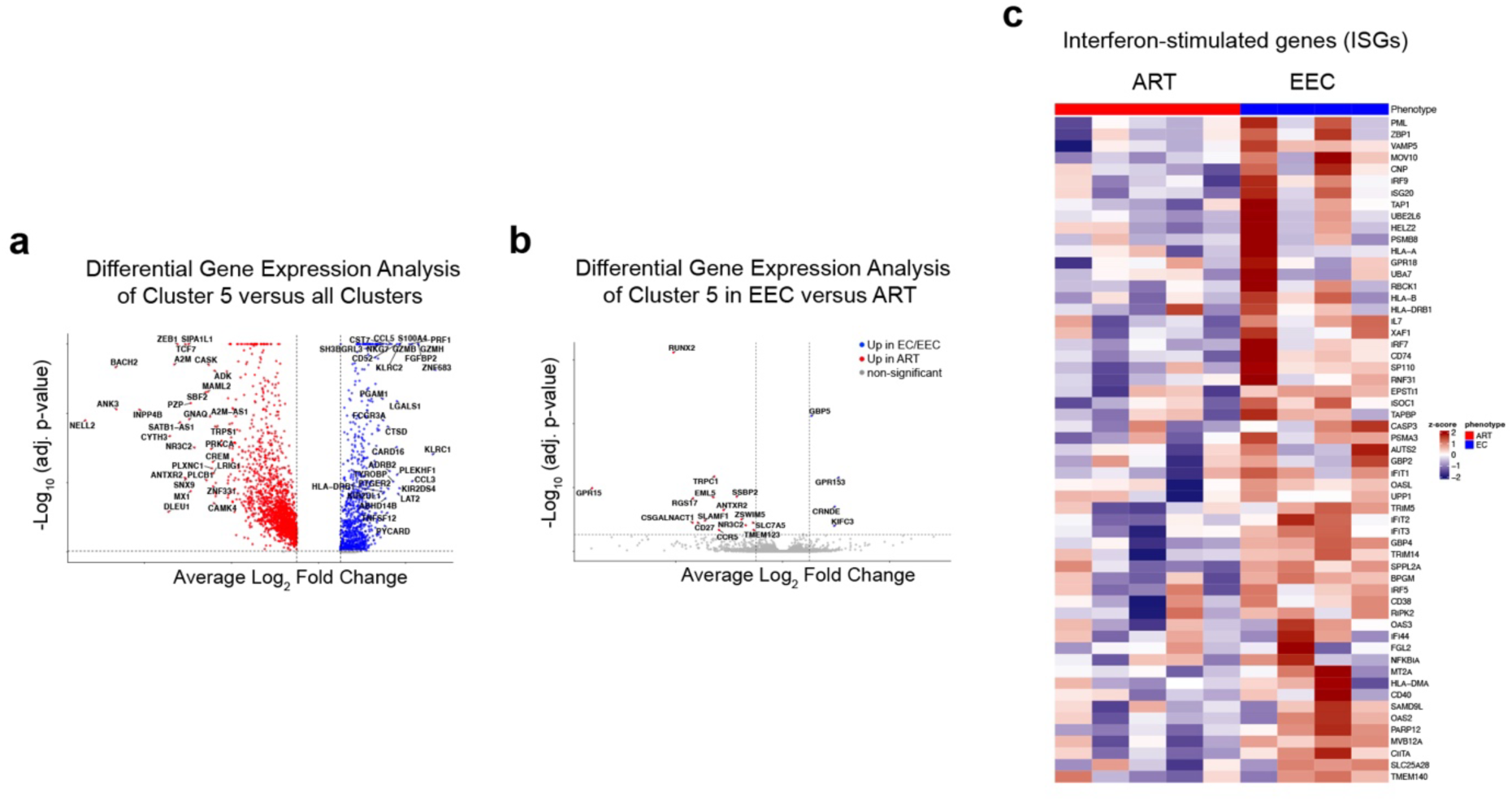
Differential gene expression and interferon-stimulated gene signature in EEC-enriched KIR⁺ CD8⁺ T cell cluster 5. **a**, Volcano plot of differential gene expression analysis comparing cluster 5 with all other KIR⁺ CD8⁺ T cell clusters, identifying canonical NK cell transcripts and cytotoxic effector genes among the most highly upregulated genes in cluster 5 (red, up in cluster 5). **b,** Volcano plot of pseudobulk differential gene expression within cluster 5 comparing EECs versus ART-suppressed individuals. EEC-enriched genes (blue) include the interferon-inducible antiviral factor *GBP5*. ART-enriched genes (red) include the T cell differentiation transcription factor *RUNX2*, the co-stimulatory and memory markers *SLAMF1* and *CD27*, and the chemokine receptor *CCR5*. **c,** Heatmap showing z-scored pseudobulk expression of leading-edge interferon-stimulated genes (ISGs) from IFN-α and IFN-γ GSEA pathways in cluster 5, displayed per donor (ART versus EEC). The ISG signature is consistently upregulated across EEC donors relative to ART donors and defines the interferon-response leading-edge module score used in Fig. 3j.

**Extended Data Figure 8.**
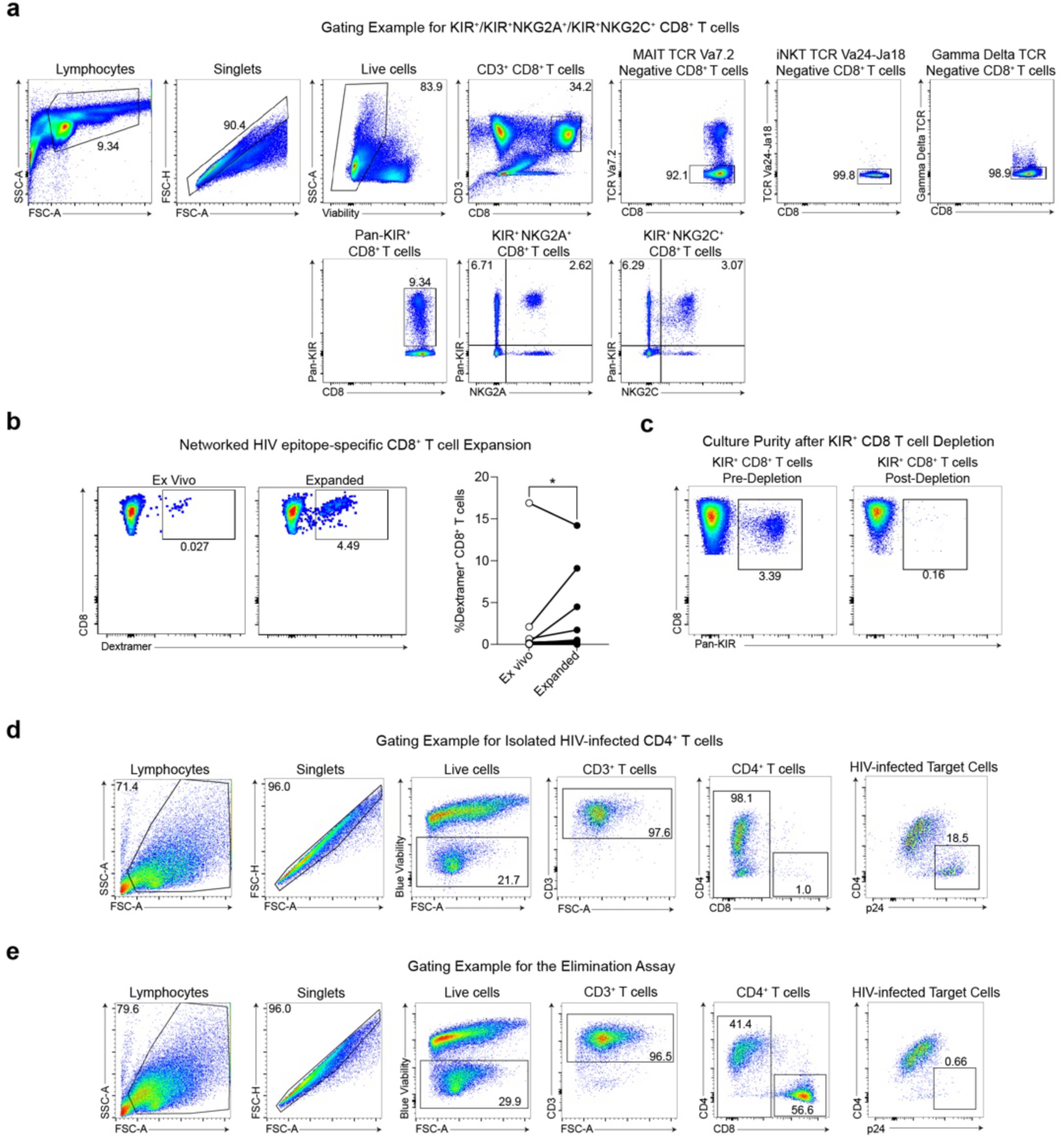
Gating strategies and quality controls for KIR⁺ CD8⁺ T cell flow cytometry and the autologous HIV elimination assay. **a**, Representative gating strategy for identification of pan-KIR⁺, KIR⁺NKG2A⁺, and KIR⁺NKG2C⁺ CD8⁺ T cell populations. Following exclusion of MAIT (TCR Vα7.2⁺), iNKT (TCR Vα24-Jα18⁺), and γδ TCR⁺ cells, pan-KIR⁺ CD8⁺ T cells are identified (6.71% in the representative donor) and subsequently gated for dual KIR⁺NKG2A⁺ (2.62%) and dual KIR⁺NKG2C⁺ (3.07%) populations. **b,** Networked HIV epitope-specific CD8⁺ T cell expansion confirmed by dextramer staining. Left and center panels show representative flow cytometry plots of dextramer⁺ CD8⁺ T cells ex vivo (0.027%) and after 6 days of networked epitope peptide stimulation (4.49%). Right panel shows summary data across EEC donors confirming significant expansion of multimer⁺ CD8⁺ T cells following antigen stimulation (* *P* < 0.05, Wilcoxon signed-rank test). **c,** Representative flow cytometry plots showing pan-KIR⁺ CD8⁺ T cell frequencies before depletion (pre-depletion: 3.39%) and after depletion using APC-conjugated KIR antibody cocktail and anti-APC microbeads (post-depletion: 0.16%), confirming efficient KIR⁺ depletion. **d,** Representative gating strategy for identifying HIV-infected CD4⁺ target cells used in the autologous elimination assay. Sequential gates identify lymphocytes, singlets, live cells (Blue Viability Dye⁻), CD3⁺ T cells, CD4⁺CD8⁻ T cells, and HIV-infected targets identified as intracellular p24⁺ (18.5% of CD4⁺ T cells in a representative infected cell preparation). **e,** Representative gating strategy for the completed elimination assay co-culture, showing identification of residual CD4⁺ T cells and HIV-infected (p24⁺) target cells after 3-day co-culture with CD8⁺ T cell effectors. Effective CD8⁺ T cell-mediated elimination results in a marked reduction in the frequency of p24⁺ CD4⁺ T cells (0.66% in the representative well shown) compared to target-only controls.

